# Sequence-independent, site-specific incorporation of chemical modifications to generate light-activated plasmids

**DOI:** 10.1101/2023.05.26.542478

**Authors:** Khoa Chung, Michael J. Booth

## Abstract

Plasmids are ubiquitous in biology, where they are used to study gene-function relationships and intricate molecular networks, and hold potential as therapeutic devices. Developing methods to control their function will advance their application in research and may also expedite their translation to clinical settings. Light is an attractive stimulus to conditionally regulate plasmid expression as it is non-invasive, and its properties such as wavelength, intensity, and duration can be adjusted to minimise cellular toxicity and increase penetration. Herein, we have developed a method to site-specifically introduce photocages into plasmids, by resynthesizing one strand in a manner similar to Kunkel mutagenesis. Unlike alternative approaches to chemically-modify plasmids, this method is sequence independent at the site of modification and uses commercially available phosphoramidites. To generate our light-activated (LA) plasmids, photocleavable biotinylated nucleobases were introduced at specific sites across the T7 and CMV promoters on plasmids and bound to streptavidin to sterically block access. These LA-plasmids were then successfully used to control expression in both cell-free systems (T7 promoter) and mammalian cells (CMV promoter). These light-activated plasmids might be used to remotely-control cellular activity and reduce off-target toxicity for future medical use. Our simple approach to plasmid modification might also be used to introduce novel chemical moieties for advanced function.

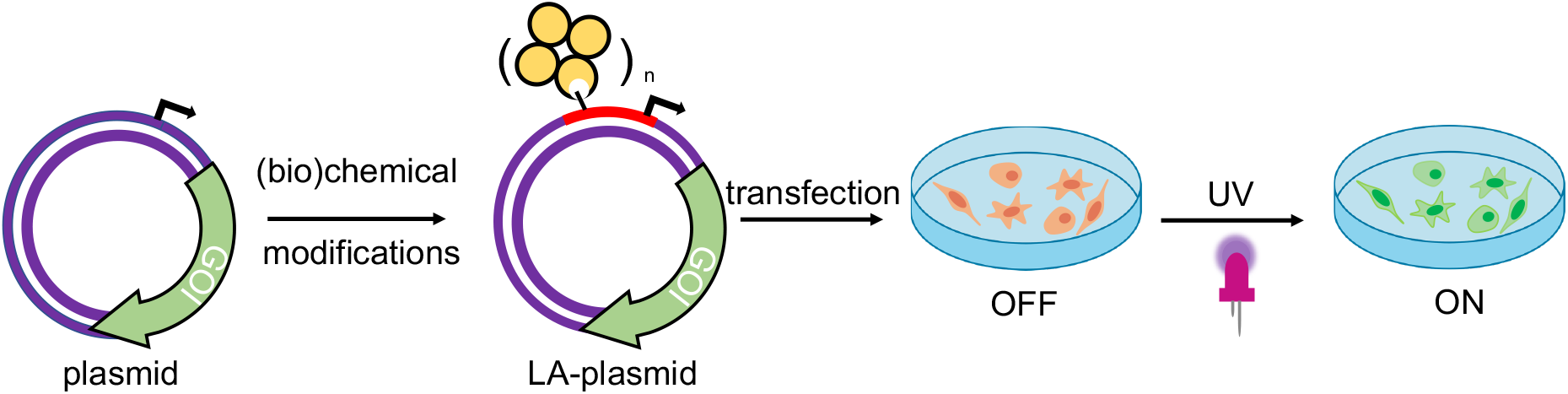

## Introduction

Biological processes are complex and occur in a tightly regulated manner. To fully understand the sophisticated interplay between one gene within a whole network, its downstream effects within a signalling cascade, or its wider physiological impacts on the cellular/tissue level, precise control over its expression is desired. Light represents an ideal means for transcriptional activation as its application is localised and its properties can be finely tuned. Photoregulation of gene expression is a burgeoning field and a multitude of light-dependent technologies are available, encompassing the entire UV-visible spectrum, to mediate photoresponses.^1^ Although natural light-responsive proteins are abundant and have shown excellent photomodulation,^2,3^ their use is limited by multi-component systems^2–9^ and lengthy optimisation.^4,10^ Alternatively, photocaging involves the installation of light-sensitive blocking groups, photocages, directly on the nucleic acids to disrupt transcription or translation. The activity is minimised/abolished until the appropriate light stimulus is applied to remove the caging groups and reactivate expression. Whilst examples of caged linear DNA^11–24^ and RNA^25–28^ are extensive, photocaged plasmids are limited in the literature.^25,29–34^ Owing to their covalently-closed nature, plasmids are resistant to exonucleases abundant within the cytoplasm and therefore they have a longer lifetime and make better templates than linear nucleic acids.^35–37^

Early examples of photocaged plasmids incorporated multiple caging groups onto the phosphate backbones in a random manner.^25,29,33,34^ Although Nagamune *et. al*. were the first to control the site of photocleavable group attachment, only one such group was present, and their method created unsealed nicks in both plasmid strands.^30^ The presence of numerous caging groups, and irregularity in their positionings across the plasmid sequence likely resulted in the leaky expression and poor activation in these initial attempts. Subsequent efforts to photocage the plasmid installed the photosensitive groups onto the Watson-Crick-Franklin (WCF) face in a site-specific manner.^31,32^ These strategies exhibited a tight OFF state, and a good ON state upon photolysis of the caging groups. However, both methods suffered from the reliance on custom phosphoramidites, which are not commercially available, to introduce the photoresponsive groups, restricting their adoption by the wider research community. Furthermore, these methods necessitated precise placements of specific enzyme-recognition or G-quadruplex forming sequences at the site of modification to enable their caging effect. Moreover, the strict presence of these sequences at or near the promoter may result in unintended changes to the promoter’s regulatory role. To circumvent these issues, we engineered a novel approach to site-specifically install photoresponsive modifications into a plasmid, by resynthesizing one strand, to govern gene expression using UV light (**Figure 1A**). By electing to block the non-WCF face of the nucleobase, all components necessary to affect plasmid resynthesis and optical control were commercially available, which will permit its routine use by any laboratory. As the short nickase recognition sites required to create these light-activated (LA) plasmids could be placed anywhere on the plasmid, this approach is general and compatible with most plasmid sequences, thus extending the approach’s accessibility. This strategy was demonstrated for two nickases with plasmids containing either the T7 or CMV promoter to control cell-free and mammalian gene expression (**Figure 1B**).

**Figure 1.**
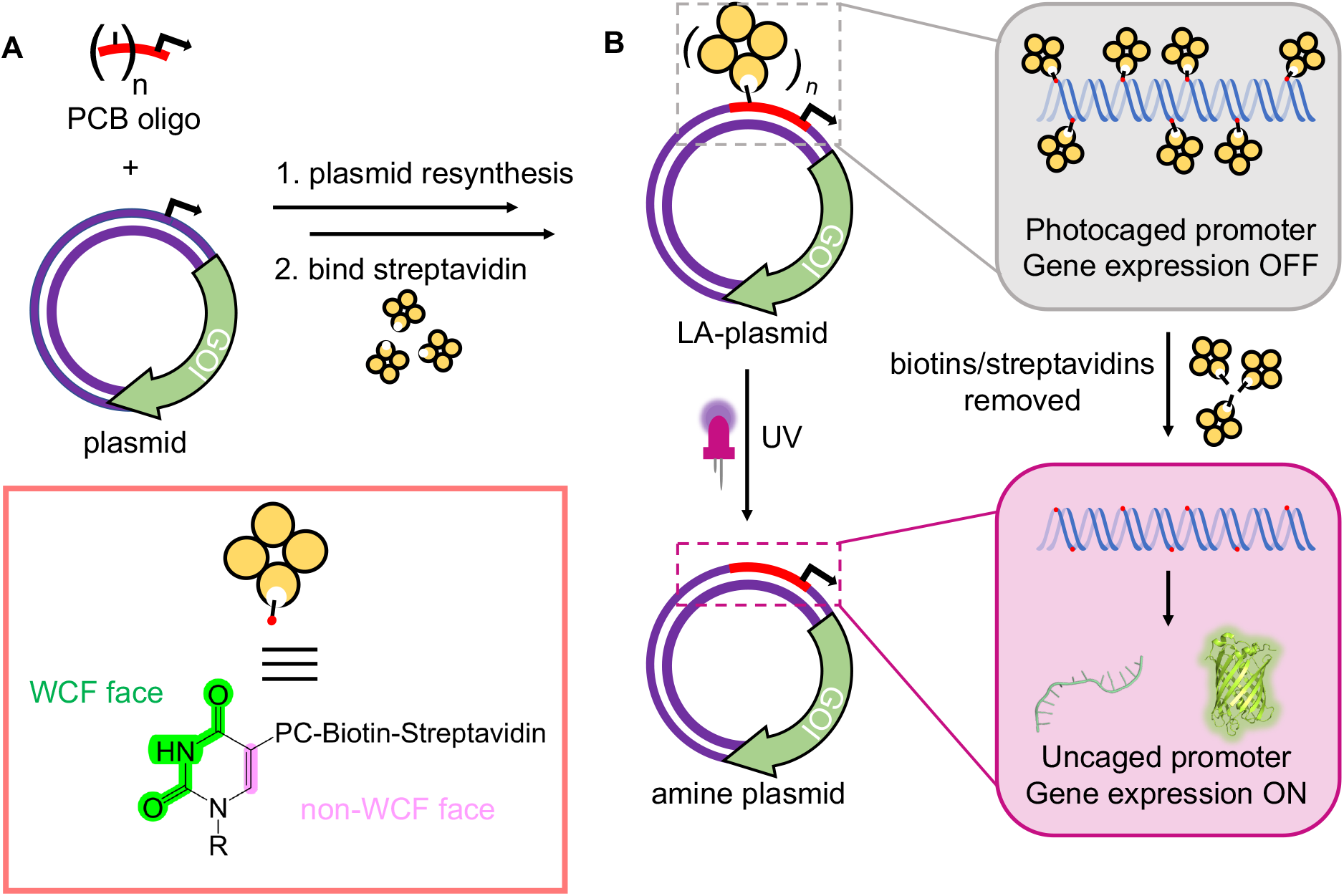
The plasmid resynthesis strategy to photocage plasmids. **A**. We have developed a method to introduce photocleavable (PC) biotin modifications site-specifically onto the non-WCF face of the nucleobases in a plasmid. **B**. Photocaging groups are incorporated across the promoter upstream of a gene of interest (GOI). Gene expression is repressed until UV illumination removes biotins/streptavidins to reactivate the plasmids.

## Results and Discussion

### Development of NEEL

To incorporate photoblocking modifications, we were inspired by Kunkel site-directed mutagenesis (SDM), which was traditionally used to introduce mutations site-specifically into DNA sequences.^38,39^ Briefly, in Kunkel SDM a plasmid, containing the sequence to be mutated, is transformed into an *E. coli* strain lacking the enzymes uracil deglycosidase (*udg*^-^) and dUTPase (*dut*^*-*^). These enzymes are involved in DNA repair by preventing dUTP insertion into bacterial chromosomes. Ultimately, bacteria produce a uracil-containing circular single-stranded DNA (cssDNA). This cssDNA serves as a template to anneal a primer bearing the desired mutations. The primer is then extended around the plasmid and ligated *in vitro*. The resultant heteroduplex comprises a mutated deoxythymine-containing strand and the original uracil-containing strand. When introduced into a *udg*^*+*^, *dut*^*+*^ *E. coli* strain, the latter strand is replaced with a thymine-containing strand.

We incorporated the *in vitro* plasmid resynthesis aspect of Kunkel SDM into our plasmid modification strategy (**Figure 2A**). Initially, the sense strand bearing the promoter and gene of interest (GOI) was reacted with a nickase to induce a single-stranded cut. The nicked strand could then be removed with an exonuclease to give a cssDNA. In a similar manner to Kunkel mutagenesis, sequential primer annealing, extension, and ligation regenerated the plasmid. This approach was termed NEEL after the four enzymatic steps: <u>N</u>icking, <u>E</u>xonuclease digestion, <u>E</u>xtension, and <u>L</u>igation.

**Figure 2.**
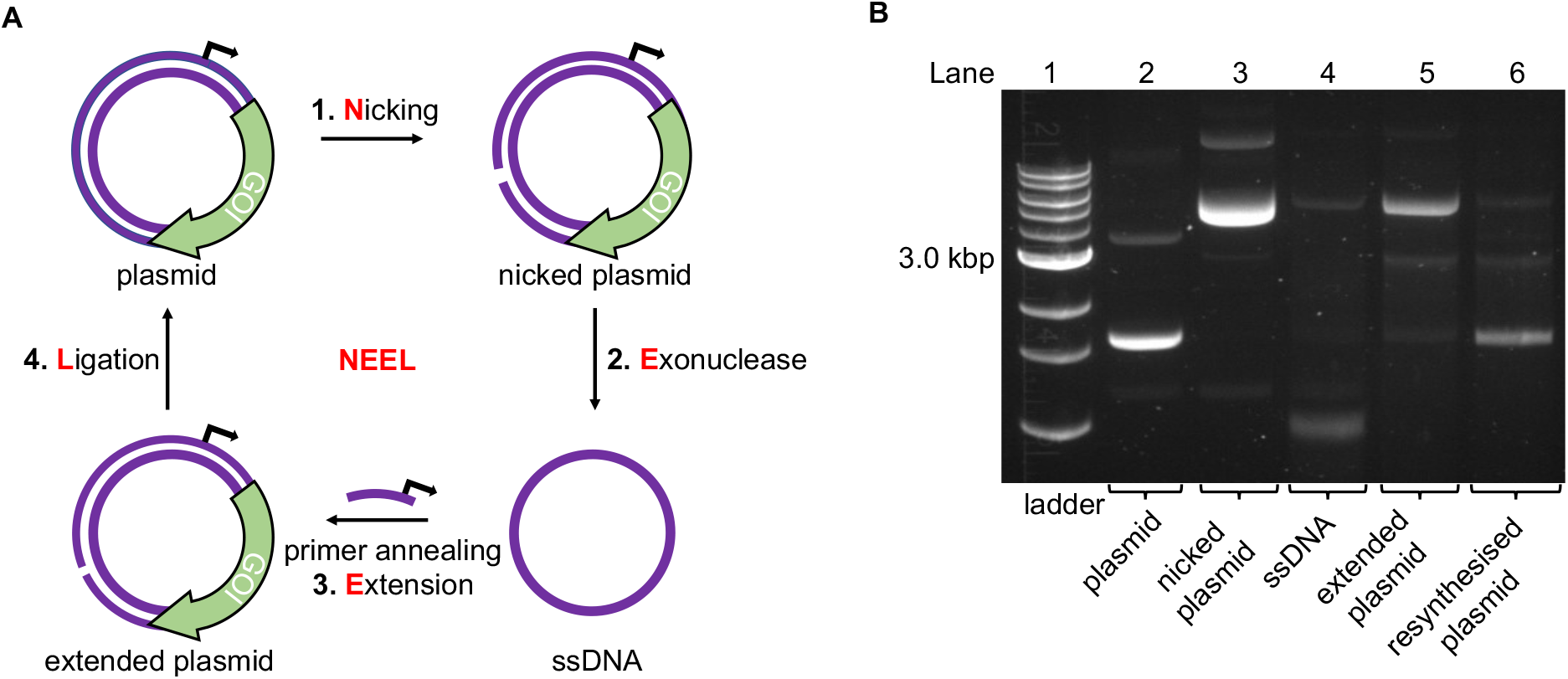
The NEEL method to resynthesise a plasmid. **A**. Overview of the enzymatic reactions in NEEL. **B**. Using NEEL, our plasmid encoding mVenus was successfully resynthesised.

Initially, a plasmid encoding for the mVenus fluorescent protein under the control of a T7 promoter was used to develop NEEL. The recognition site for the nickase Nt.BspQI was cloned downstream of the mVenus open reading frame, greater than 750 base pairs from the promoter. Theoretically, the nicking site could be positioned anywhere on the plasmid, which is an advantage of this method over a previous attempt to photocage plasmids.^31^ Following nicking and exonuclease treatment, multiple Kunkel protocols from the literature were employed to perform NEEL (**SI Figure 1**).^40–43^ It was paramount that the plasmids obtained from NEEL were ligated, as it was well known that linear DNA templates are degraded by abundant exonucleases within the cellular milieu, which hamper their transfection efficiency.^35,36^ Thus, to confirm ligation had occurred, the plasmids were assessed by agarose gel electrophoresis, with and without gyrase. Gyrase supercoils DNA and can only accept covalently-closed substrates, demonstrating the resynthesis of intact double-stranded plasmids. Without gyrase, the ligated plasmids appeared as ladder bands. However, with gyrase, these ladder bands were converted into a single band that co-migrated with the native supercoiled plasmid, confirming that ligation was successful. The non-ligated plasmid co-migrated with the nicked plasmid and remained at the same position even after reaction with gyrase.

The Kunkel protocol with the highest degree of ligation and supercoiling, and requiring a short reaction time, (Kay *et. al*.) was chosen for NEEL.^43^ We determined that the crucial determinant for NEEL success was the use of T7 DNA Polymerase (DNAP), as ligated plasmids could not be generated with other DNAPs. To verify the generality of NEEL, the dinucleotide at the ligation junction was iteratively assessed. Sixteen primers with different T_m_, GC content, and lengths, annealing at different positions across the plasmid were used for NEEL (**Supplementary Table 1** and **SI Figure 2A** and **2B**). These primers were also chosen to include all the possible dinucleotide combinations at the ligation junction. Ligation was successful regardless of primer features, annealing location, and the dinucleotide identities at the ligation junction. Having confirmed ligation was robust, gels were pre-stained hereafter as the resynthesised plasmid bands were more intense and easier to distinguish as they co-migrated with the native plasmid due to GelRed-DNA interactions (**Figure 2B**).

To optimise the plasmid resynthesis, we wanted to remove any native plasmids that were not nicked or removed by exonuclease, which would contribute to an increase in the OFF-state when using photocages. To do this we used DpnI, a nuclease that preferentially cleaves dam-methylated DNA over non-methylated substrates.^44^ NEEL resynthesised plasmids were hemi-methylated, as they contained a parental methylated strand and a synthetic unmethylated strand. Although DpnI can cleave hemi-methylated DNA, its cleavage activity is significantly lower.^45^ Therefore, mild DpnI conditions were optimised on the native plasmid, to minimise degradation of the resynthesised plasmid (**SI Figure 3A**). Compared to the supplier’s recommended conditions, plasmid degradation could be achieved using a much lower enzyme concentration and shorter reaction time. When tested with the resynthesised plasmid, faint bands denoting degradation of the methylated (major) and hemi-methylated plasmids (minor) were detected, but the resynthesised plasmid remained the most intense band (**SI Figure 3B**).

**Figure 3.**
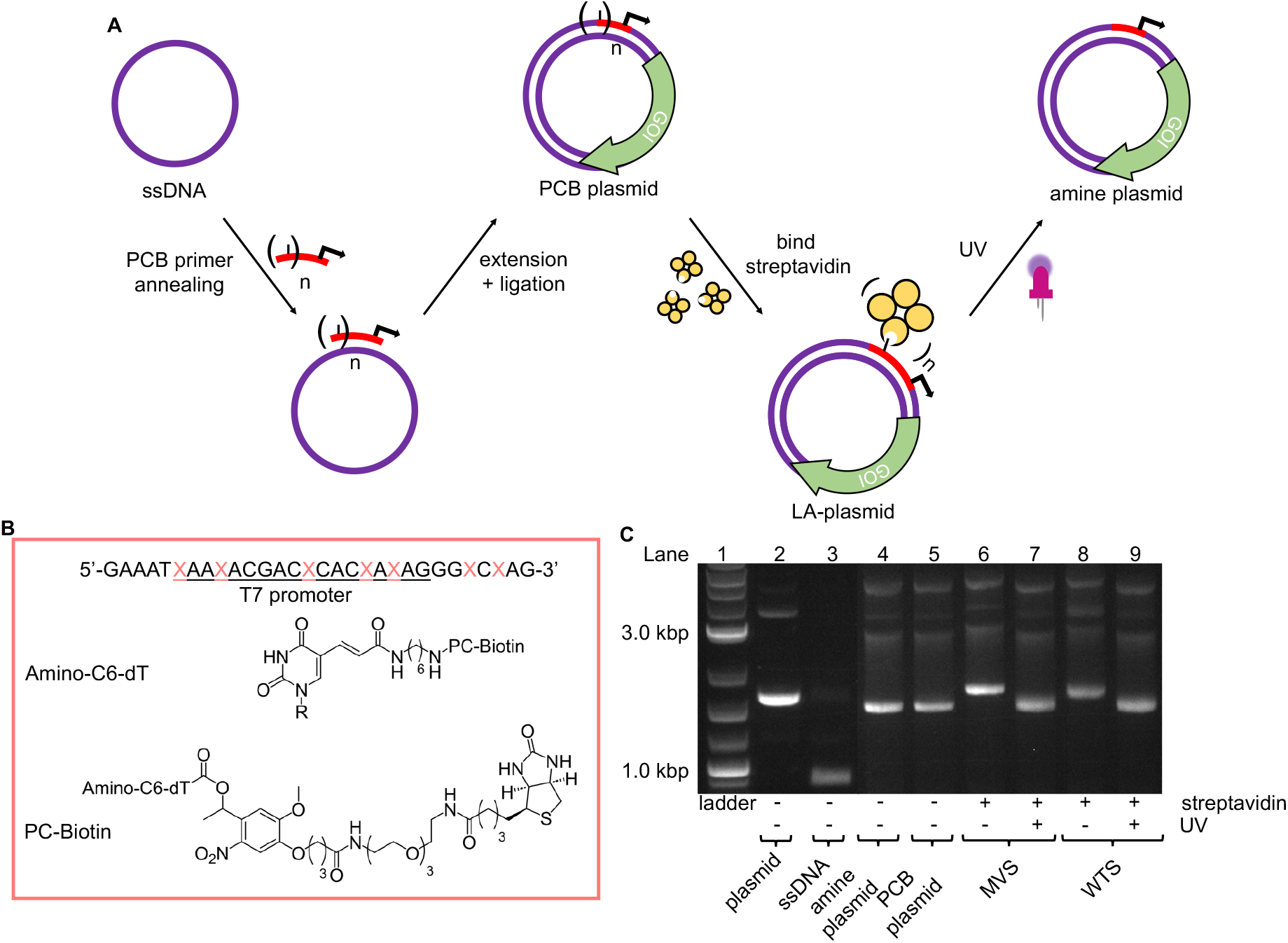
LA-plasmids through photocaging the T7 promoter. **A**. Overview of strategy to prepare LA-plasmids. **B**. The T7 amine primer where the T7 promoter sequence is underlined, and X is the modified base. Structures of the amine and PCB modifications are also shown. **C**. T7 LA-plasmids and their photouncaging.

### Photocaging the T7 Promoter

With a robust method to resynthesise the plasmid in place, photocaging groups were then introduced to the sense strand of the promoter to control the expression of a downstream gene. To this end, the light-activated DNA (LA-DNA) platform previously developed in our group was used as a starting point.^14^ In this approach, an oligo containing seven amines across the T7 promoter sequence is reacted with a UV-photocleavable 2-nitrobenzyl biotin N-hydroxysuccinimide (NHS) ester. The photocleavable biotin (PCB) oligo was used then as a primer in PCR to create a linear PCB DNA template, with the PCB T7 promoter upstream of a GOI. Binding streptavidin blocks T7 RNA Polymerase (RNAP) from accessing the promoter, thereby inhibiting transcription. Conditional gene expression is restored upon UV-triggered removal of the biotins/streptavidins.

Analogous to the above linear LA-DNA templates, LA-plasmids were prepared by resynthesising the plasmid using a PCB T7 promoter oligo, followed by binding of streptavidin (**Figure 3A**). The T7 amine primer was reacted with PCB-NHS ester, purified by HPLC, and analysed by LC-MS and denaturing PAGE (**Figure 3B**, and **SI Figure 4A** and **4B**). Using mild UV conditions (∼1.06 mW cm^-2^ for 4 mins), multiple bands, representing successive removal of PCB groups, were observed. We then demonstrated that the PCB remained intact when subjected to annealing conditions, and its association with streptavidin was not disrupted (**SI Figure 5**).

Excitingly, plasmid resynthesis using the amine or PCB T7 oligo gave amine and PCB plasmids, respectively, which illustrated T7 DNAP’s high tolerance for base modifications (**Figure 3C**). The latter plasmid was bound to either monovalent or wildtype tetravalent streptavidin (MVS and WTS, respectively) to give LA-plasmids. These could be observed in the gel by a decrease in electrophoretic mobility. Interestingly, little interplasmid crosslinking was observed using the WTS, which may be due to the concentrations involved and/or the favourability of intrastrand crosslinks (**SI Figure 6**). As expected, UV illumination removed biotins/streptavidins to restore the amine plasmids. Plasmids resynthesised using a regular primer without amino modifications did not show any changes using similar conditions (**SI Figure 7**). This confirmed the requirement of both the amine and PCB groups to generate LA-plasmids.

These T7 promoter LA-plasmids were then applied in cell-free gene expression systems to determine their photomodulation efficiency. *In vitro* transcription (IVT) with the T7 RNAP showed that the cssDNA was not transcriptionally active (**Figure 4A**). Whereas the resynthesised plasmid without modifications or containing amine and PCB modifications all produced transcripts at the expected size, indicating that plasmids produced using the NEEL method were functionally active, and the T7 RNAP was tolerant to minor base modifications at the T7 promoter. However, when bulky streptavidin was appended, the T7 RNAP was unable to associate with its cognate promoter, and transcription was inhibited. Following UV illumination, the steric block was released and transcription from the plasmid was restored. Cell-free protein synthesis (CFPS), using the commercially available NEBExpress system, revealed this control was also measurable at the translational level (**Figure 4B** and **SI Figure 8**). Expression from caged plasmids was abolished (96% for both the MVS and WTS, compared to the ‘amine plasmid -UV’ condition). Upon UV-induced cleavage of the PCB moieties, the fluorescence increased by 19-fold for plasmids caged with either MVS or WTS. The uncaged MVS and WTS LA-plasmids obtained 77% and 67% relative to the ‘amine plasmid +UV’ condition (although non-significantly different), respectively. These data clearly showed that NEEL produced T7 promoter LA-plasmids that tightly controlled gene expression.

**Figure 4.**
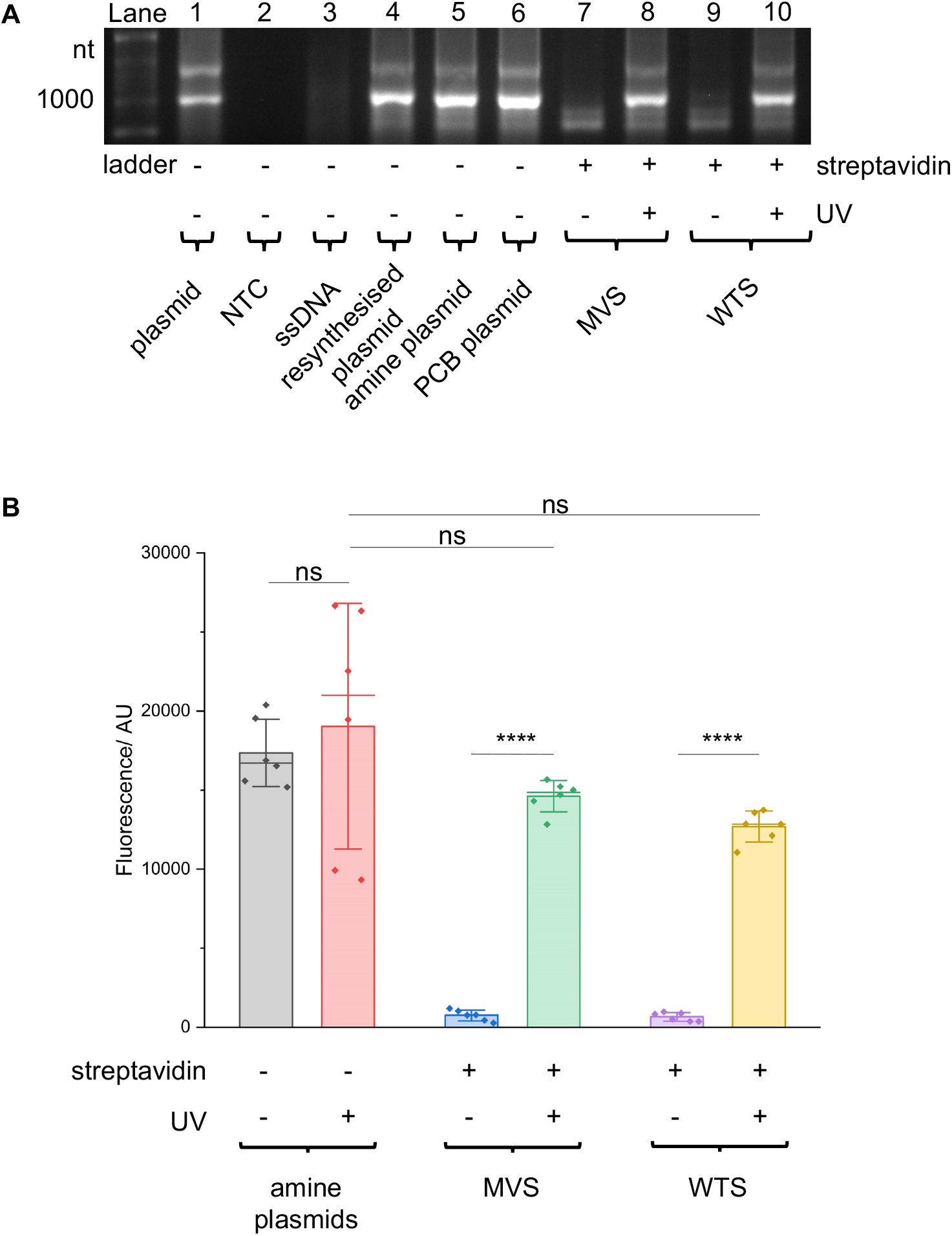
T7 promoter photocaged LA-plasmid activity in cell-free gene expression. Plasmids resynthesised using NEEL were active in **A**. IVT and **B**. CFPS. When these plasmids were photocaged, the resultant LA-plasmids showed UV-dependent gene expression. In (**B**) fluorescence was background-subtracted from NTC (*n =* 3). All other conditions were biological duplicates of technical triplicates (*n* = 6). Error bars showed standard deviations. Two-tailed, unpaired Student’s t-test was performed. ns = non-significant (P > 0.05); ****P ≤ 0.0001.

### Photocaging the CMV Promoter

To demonstrate the generality of the NEEL protocol and LA-plasmid technology, we applied these approaches to exert conditional control of the cytomegalovirus (CMV) promoter, a widely used promoter sequence in eukaryotic expression systems. This promoter has been extensively studied and its core promoter elements are well-characterised, thus providing a good platform to illustrate our caging strategy in mammalian cells. Transcription initiation commences with the formation of the preinitiation complex (PIC) at the promoter.^46–50^ This process begins with the recognition of the TATA-box sequence ∼30 bp upstream of the transcription start site (TSS) by its cognate partner, the TATA-box binding protein (TBP), a subunit of the general transcription factor (GTF) TFIID.^51^ The initiator element (Inr) is a 6-7 bp motif that covers the TSS and directs where transcription begins.^52^ After TFIID binding, TBP recruits TFIIB whereby the latter can bind to the B-recognition elements immediately upstream and downstream of the TATA-box (BRE^u^ and BRE^d^, respectively).^53^ These early interactions engage other components of the PIC such as RNAP II and other GTFs.^54^

It has been shown *in vivo* that RNAP II transcription required all the GTF components.^46^ Their proximal positions relative to one another, as well as the availability of the full and partial consensus sequences (**Table 1**) permitted us to examine these sites to exert light-mediated gene expression. The Nt.BbvCI recognition site was cloned greater than 1,000 base pairs away from the CMV promoter, in a mammalian expression plasmid encoding the green fluorescent reporter protein mNeonGreen. This plasmid was then subjected to the LA-plasmid caging strategy.

**Table 1.** Comparison between plasmid core promoter elements and the corresponding consensus sequences in the CMV promoter.

| Element | Consensus sequence | Plasmid sequence | Matching sequence |
| --- | --- | --- | --- |
| TATA box | TATAWAWR<sup>55</sup> | <u>TATATAAG</u> | 8/8 |
| Inr | YYANWYY<sup>56,57</sup> | <u>CCATTCT</u> | 7/7 |
| BRE<sup>u</sup> | SSRCGCC<sup>58</sup> | <u>GGAGGTC</u> | 5/7 |
| BRE<sup>d</sup> | RTDKKKK<sup>59</sup> | <u>CAATGCT</u> | 4/7 |

(W) A or T; (R) A or G; (S) G or C; (D) A, G, or T; (K) G or T; (Y) C or T; (N) A, C, G, or T. Plasmid sequence with underlined bases matched those of the consensus sequence.

A CMV promoter oligo containing nine amino modifications in place of thymines across the regulatory regions was used to prepare the PCB oligo (**Figure 5A**). Once again, LC-MS and denaturing PAGE were used to confirm PCB attachment (**SI Figure 9A** and **9B**). The UV parameters were also increased (9.88 mW cm^-2^ for 10 mins) to ensure the full removal of PCB groups and match the UV conditions previously used with mammalian cells in the literature.^21^ As has also been demonstrated by others, these UV settings were not detrimental to cell viability (**SI Figure 10**).

**Figure 5.**
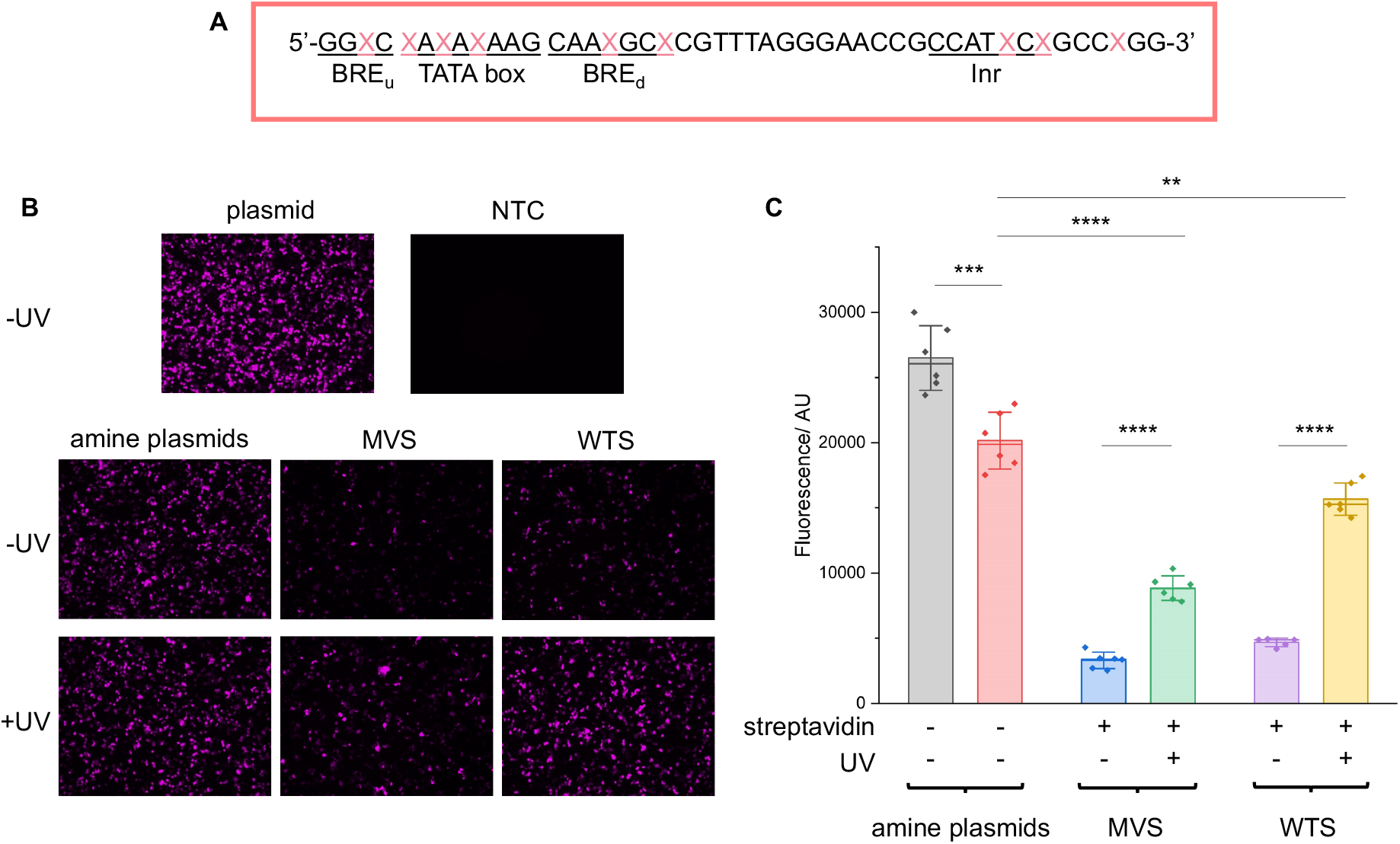
CMV promoter photocaged LA-plasmid activity in mammalian cells. **A**. The CMV amine primer, where the CMV core promoter elements are underlined and X is the modified base. The NEEL plasmids were transfected into HEK293T cells and assessed for their activity with and without UV irradiation. Expression was measured through **B**. Fluorescence microscopy; images were representative of the respective conditions. **C**. Fluorescence plate reader; the fluorescence was background-subtracted from NTC. Error bars showed standard deviations. Two-tailed, unpaired Student’s t-test was performed. ns = non-significant (P > 0.05); **P ≤ 0.01; ***0.0001 ≤ P ≤ 0.001 and ****P ≤ 0.0001. For (**B**), ‘plasmid’ and ‘NTC’ conditions were technical triplicates (*n =* 3), and all other conditions in (**B**) and (**C**) were biological duplicates of technical triplicates (*n* = 6) except for the ‘WTS-UV’ condition (*n =* 5 due to sample loss).

LA-plasmids containing either 9 MVS or WTS were prepared through plasmid resynthesis using the PCB CMV promoter oligo and analysed by agarose gel electrophoresis, demonstrating protein binding and release with UV illumination (**SI Figure 11**). These plasmids were transfected into HEK293T cells and expression was assessed with and without illumination. In the absence of UV, transcription, measured by RT-qPCR, from the LA-plasmids was reduced by 82 and 80% relative to the ‘amine plasmid -UV’ condition for MVS and WTS LA-plasmids, respectively (**SI Figure 12A**). Upon UV uncaging, transcription was increased by ∼2 and 4.5 times for the MVS and WTS-caged plasmids, respectively. The latter represented a 100% recovery when compared to the ‘amine +UV’ control. Similar results were recapitulated at the translational level. Notably, epifluorescence microscopy confirmed that both LA-plasmid types were repressed (**Figure 5B**). UV application had no discernible effect on the amine plasmid but gave a significant increase in fluorescence for the LA-plasmids. To quantify the change in protein levels a fluorescence plate reader was used (**Figure 5C** and **SI Figure 12B**). Protein expression was diminished by 88% and 82% relative to the ‘amine plasmid -UV’ condition, and the increase in expression following illumination was 2.7 and 3.3-fold for the MVS and WTS LA-plasmids. Relative to the amine plasmid control, these numbers translated to a 44% and 78% recovery. This demonstrated light-activated control of expression in mammalian cells using commercially available materials.

## Conclusion

We have developed a simple and accessible approach to photoregulate plasmid expression in cell-free systems and mammalian cells. Inspired by the classical Kunkel mutagenesis method, NEEL was devised to resynthesise plasmids through four concerted enzymatic reactions using the nickase, exonuclease, polymerase, and ligase enzymes. Unlike previous approaches that relied on random, uncontrolled, and excessive caging of the phosphate backbones,^25,29,33,34^ our method site-specifically installed the caging modifications by using a primer that contained photocleavable biotins (PCB). Additionally, our method allows for the placement of a nickase site anywhere on the plasmid, unlike previous approaches that required the incorporation of sequence-specific motifs around the promoter site.^31,32^ We used our LA-plasmid technology to regulate expression from the T7 promoter in cell-free protein synthesis and the CMV promoter in mammalian cells. The incorporation of streptavidin proteins across the promoters provided a tight caging of activity that could be efficiently reconstituted following UV illumination.

Owing to the availability of nickases with diverse recognition sequences, we expect the LA-plasmid caging approach to be generalisable to most plasmid sequences. Furthermore, with the flexibility of placement of the nickase site, the commercial availability of all reagents, and the simplicity of the workflow, we envisage this strategy could be easily adopted by non-chemistry-based laboratories to photoregulate plasmid expression. This would be especially relevant for studies in developmental biology^26,27^ and population dynamics.^60^

The application of LA-plasmids in cell-free systems would allow the use of cheaper lysate systems that cannot accommodate linear DNA templates due to the presence of nucleases. Finally, the multivalency of streptavidin provides opportunities to incorporate other functionalities onto the plasmid. An interesting avenue is cell-penetrating moieties that would permit light-activated, self-delivered plasmid constructs. This is made possible by the availability of diverse biotinylated reagents such as transferrin and albumin that have been shown to enable gene delivery.^61,62^ The recent approvals of plasmid-based gene therapies^63–65^ and vaccines^66,67^ in humans signalled a renewed interest in plasmids as therapeutic agents. Thus, the ability to precisely control their expression with a non-intrusive external stimulus such as light is timely and could limit off-target expression and unexpected outcomes.

## Materials and Methods

Unless otherwise stated, all enzymes were purchased from New England Biolabs (NEB). Seakem® LE Agarose was purchased from Lonza. XL-10 Gold competent *Escherichia coli* (*E. coli*) cells and SOC Outgrowth Medium were purchased from NEB. ATP, SuperScript IV VILO Master Mix with ezDNase, Fast SYBR Green Master Mix, Lipofectamine 3000, Fluorobrite DMEM, Opti-MEM reduced serum medium, Genejet PCR Purification Kit were purchased from Thermo Fischer Scientific. Non-modified primers were purchased from Integrated DNA Technologies. Amine primers were purchased from ATD-Bio or Merck/Sigma. PCB-NHS ester was purchased from Click Chemistry Tools or Merck/Sigma and stored in dry dimethylformamide (DMF) at -80 ⁰C. HPLC solvents, trypsin-EDTA, FBS, Amicon 3K columns, 3X GelRed, and TAE Buffer were purchased from Merck/Sigma. TEMED was purchased from Bio-Rad. Gel extraction, maxiprep, miniprep, and RNeasy Mini kits were purchased from Qiagen. CellTiter-Glo Luminescent Cell Viability Assay was purchased from Promega. Pico488 dsDNA Quantification Reagent and the dsDNA quantitative standard (100 ng/µL) in TE Buffer were purchased from Lumiprobe. HEK293T cell line was a gift from Dr. Haiping Tang (Prof. Dame Carol Robinson’s group). The mVenus plasmid (containing the T7 promoter) was available and previously prepared by Booth *et. al*.^14^ The pCS2-mNG-C (the CMV mNG plasmid) was purchased from Addgene (plasmid #128144).

### Agarose Gel Electrophoresis

100-150 ng DNA was analysed on a 1.0% pre-stained (1X GelRed) agarose gel. The gel was run in 1X TAE Buffer at 130 V for 55 mins. Bands were imaged using a BioRad Molecular Imager® Geldoc™ XR+ gel imager and analysed using ImageLab 6.1.

### Preparing the T7 mV and CMV mNG Plasmids

Homologous recombination was used to insert the BbvCI and BspQI recognition sites into the T7 mVenus plasmid to create mV_nicks, and the BbvCI recognition site into the CMV mNG plasmid to create mNG_BbvCI (**Supplementary Table 2**). For the T7 mVenus plasmid, the plasmid was digested to form a linear fragment with BamHI before PCR amplification. For the CMV mNG plasmid, the circular plasmid was used directly for PCR. Separate PCR reactions were carried out to amplify the backbone and insert. The reactions were performed in 0.2 mL PCR tubes and contained the native or digested plasmid template (1-3 ng), the forward and reverse primers (1 µL, 10 µM, **Supplementary Tables 3-5**), Phusion High-Fidelity PCR Master Mix with HF Buffer (10 µL, 2X), and made up to 20 µL with MilliQ water.

Reactions were cycled on a Peqlab peqSTAR 96X Universal thermal cycler using the following conditions: initial denaturation at 98 °C for 30 seconds; 30 or 35 cycles [denaturation at 98 °C for 10 seconds, annealing at either 55 °C (mV_nicks) or 68 °C (mNG_BbvCI) for 30 seconds, and extension at 72 °C for either 1.5 minutes (backbone and insert of mV_nicks) or 30 seconds (mNG_BbvCI insert) and 2.5 minutes (mNG_BbvCI backbone)]. A final extension at 72 °C for 10 minutes was performed. DpnI (0.5 µL, 20 U/µL) was added and reactions were incubated at 37 °C for 30 minutes and heat-deactivated at 80 °C for 20 minutes to remove the templates. The reactions were purified with a Genejet PCR column according to the manufacturer’s protocol. Reactions were analysed with gel electrophoresis to confirm product bands.

A 7:1 insert:backbone ratio was used for homologous recombination and calculated according to NEBioCalculator. The backbone (100 ng) and insert were combined in a 1.5 mL Eppendorf tube. XL-10 Gold competent *E. coli* cells (50 µL) were added to the reaction. The tube was kept on ice for 30 minutes and heat-shocked at 42 °C for 30 seconds. SOC Outgrowth Medium (500 µL) was added and the tube was incubated at 37 °C with shaking at 225 rpm for 30 minutes. The tube was kept on ice for 2 minutes and 200 µL of the transformed bacteria were plated on LB agar plates containing ampicillin (100 µg/mL) at 37 °C for 14-16 hours. Colonies were then picked and grown in 50-mL Falcon tubes containing 5 mL LB media with ampicillin (100 µg/mL) at 37 °C and shaking at 225 rpm for 14-16 hours. 300 µL of the overnight cultures were combined with glycerol (300 µL, 50% v/v, filtered). The glycerol stocks were kept at -80 °C. The rest of the cultures were miniprepped using Qiagen’s miniprep kit according to the manufacturer’s protocol and eluted in their Elution Buffer (50 µL) and stored at -20 °C. Sanger sequencing was performed by SourceBioScience. After the sequence was confirmed, bulk plasmid was obtained by inoculating 2 × 100 mL LB cultures with the desired glycerol stock. The cultures were grown as above and purified using Qiagen’s Maxiprep kit and performed according to the manufacturer’s protocol. The plasmids were resuspended in Tris (1 mL, 10 mM, pH 8.0) and stored at -20 °C.

### Nicking the T7 mV and CMV mNG Plasmids

To a 1.5 mL Eppendorf tube, the mV_nicks plasmid (75 µg), Nt.BspQI (62.6 µL, 10 U/µL), and NEBuffer 3.1 (62.6 µL, 10X) were added and made up to 625 µL with MilliQ water. The reactions were incubated on the 1.5 mL Thermomixer C (Eppendorf) with shaking at 600 rpm at 50 °C for 2 hours and heat-deactivated at 80 °C for 25 minutes.

The nicking reactions for the mNG_BbvCI plasmid were performed analogously but instead used the Nt.BbvCI nickase (62.6 µL, 10 U/µL) and CutSmart Buffer (62.6 µL, 10X). The reactions were incubated on the Thermomixer C with shaking at 600 rpm at 37 °C for 2 hours and heat-deactivated as above. These reactions were then purified using Geneject PCR columns according to the manufacturer’s protocols and eluted in their Elution Buffer (50 µL). The concentrations were measured using the Nanodrop ND-1000 Spectrophotometer and stored at -20 °C.

### Preparing cssDNA Templates

The total reaction volumes varied and depended on the amount of nicked plasmid used. For example, when 60 µg dsDNA template is used, the total reaction volume would be 500 µL. The reactions were performed in 1.5 mL Eppendorf tubes and consisted of the plasmid (final concentration = 120.0 ng/µL), Lambda Exonuclease (final concentration = 1.0 U/µL), Lambda Exonuclease Reaction Buffer (final concentration = 1X), and made up to the desired volumes with MilliQ water. The reactions were incubated on a 1.5 mL Thermomixer C with shaking at 600 rpm at 37 °C for 3 hours, and heat-deactivated at 75 ⁰C for 20 minutes. The reactions were purified using Genejet PCR columns according to the manufacturer’s protocol and eluted in their Elution Buffer (50 µL). The concentrations were measured using the Nanodrop and stored at -20 °C.

### PCB-NHS Ester Reaction with the Amine Oligos

Sodium bicarbonate (1 M) was prepared fresh and used within 1 week. The reactions were performed in 1.5 mL Eppendorf tubes and consisted of the amine oligo (20 µL, 100 mM), sodium bicarbonate (20 µL, 1 M, pH ∼8.5), PCB-NHS ester (100 µL, 10 mM), and made up to 200 µL with MilliQ water. The reactions were incubated with shaking at 600 rpm at 20 °C overnight.

Tris (200 µL, 10 mM, pH 8.0) was added and the solutions were purified on the Amicon 3 K 0.5 mL column by centrifugation at 14,000 g for 15 minutes at room temperature. This was performed 3 times in total, each time decanting the eluate and making up the solution to 500 µL with MilliQ water. The concentrate was eluted by inverting the column and centrifuging at 1,000 g for 2 minutes into a collection tube. This was repeated once more to maximise yield. The solution was made up to 300 µL with MilliQ water.

### Purification of the PCB Oligos

The PCB oligos were HPLC purified. Lines B/C/D were: MilliQ water, acetonitrile, and triethylammonium bicarbonate buffer (TEAB, 100 mM, pH 8.5), respectively. Line D was kept constant at 10%. Solvent percentages referred to line C, the organic phase. The flow rate was kept at 1.5 mL/minute. Separations were UV-monitored at wavelengths 260, 280, 310 and 360 nm.

The oligo (100 µL, ∼6.67 µM) was injected into an Agilent Polaris 5 C18-A (150 × 4.6 mm) analytical column heated to 50 °C. The column was equilibrated at 5% for 20 minutes. A gradient of 5-30% over 27 minutes was used to purify the oligo. The column was washed with 90% acetonitrile from 27.01 until 30 minutes. The organic phase was returned to 5% at 30.01 minutes and kept until 33 minutes for column equilibration. Three injections were made, and the fractions were collected according to **Supplementary Table 6**.

The fractions were then combined and lyophilised on the Christ Alpha 2-4 LDplus to dryness. The reactions were then resuspended in MilliQ water (500 µL) and purified using Amicon 3 K 0.5 mL columns as previously. The eluate was made up to 1 mL with MilliQ water and lyophilised to dryness. The PCB oligo was resuspended in Tris (50 µL, 10 mM, pH 8.0) and the concentration was measured using the Nanodrop.

### Denaturing PAGE of Amine and PCB Oligos

A 40% w/v polyacrylamide (PAA) stock solution was prepared by dissolving 12 g PAA in MilliQ water and made up to 30 mL with MilliQ water, wrapped in tin foil and stored at 4 °C. A 10% w/v ammonium persulphate (APS) stock solution was prepared by dissolving APS (1 g) in MilliQ water and the solution was made up to 10 mL with MilliQ water and 1 mL aliquots were kept at -20 °C. A 1 mL 2X loading dye was prepared and consisted of formamide (95% v/v), sodium dodecyl sulphate (0.02% w/v), xylene-cyanol (0.01% w/v), bromophenol blue (0.02% w/v), and EDTA (10 µL, 1 M, pH 8.0).

The amine and PCB oligos were analysed on a denaturing PAGE. The 16% PAA gel solution was prepared in a 15-mL Falcon tube and consisted of TBE Buffer (0.5 mL, 10X), PAA (2.0 mL, 40% w/v stock), GelRed (1 mL, 3X), urea (2.10 g). The solution was vigorously vortexed and sonicated and degassed at 35 °C for 10 minutes. The solution was vortexed again to ensure all the urea dissolved. APS (40 µL, 10% w/v) and tetramethylethylenediamine (2.5 µL) were added to the PAA solution, inverted several times to mix, and quickly loaded onto a pre-assembled glass plate and allowed to set.

The oligo samples consisted of the nucleic acid (200 ng) and a homemade loading dye (5 µL, 2X), and the solutions were made up to 10 µL with MilliQ water. PCB oligo receiving UV was prepared without the dye, irradiated with UV and the dye was added as above. The solutions were run in 1X TBE Buffer at 200 V for 1 hour. The gel was soaked in GelRed (3X) for 5 minutes and imaged on a BioRad Molecular Imager® Geldoc™ XR+ gel imager and analysed using ImageLab 6.1.

### LC-MS of PCB Oligos

Line A was: 8.6 mM Et_3_N, 200 mM HFIP in 5% MeOH/H_2_O (v/v). Line B was: 20% buffer A in MeOH. Solvent percentages referred to line B.

Samples (10 µL, 1 µM) were prepared and injected onto a Waters Acquity UPLC BEH C18 column (130 Å, 1.7 µm, 2.1 mm x 50 mm). Samples were analysed over 8 mins with a 0-70% gradient. Masses were recorded on a Waters Xevo G2 QTOF ESI-UPLC-MS system. Analysis was carried out using MassLynx v4.1.

### Phosphorylation of Oligos

The reactions were performed in 0.2 mL PCR tubes and consisted of the non-modified or amine or PCB oligo (60 pmol, **Supplementary Table 3**), T4 Polynucleotide Kinase (2 µL, 10 U/µL), T4 Polynucleotide Kinase Reaction Buffer (4 µL, 10X), ATP (0.4 µL, 100 mM) and made up to 40 µL with MilliQ water. The reactions were incubated on the thermal cycler at 37 °C for 1 hour, heat-deactivated at 80 °C for 20 minutes, and stored at -20 °C.

### Annealing Oligos to the cssDNA Template

The total reaction volumes varied and depended on the amount of cssDNA used. For example, when 10 µg cssDNA template is used, the total reaction volume would be 779 µL. The molar mass of the cssDNA template was calculated based on the template length and assuming each base had an average mass of 330. The molar amount could then be deduced.

The annealing reactions were performed in 1.5 mL Eppendorf tubes and consisted of the cssDNA template (final concentration = 12.84 ng/uL), the phosphorylated primer (4 molar equivalents relative to the cssDNA template), magnesium chloride (final concentration = 10 mM), Tris (pH 7.4, final concentration = 50 mM), and made up to the desired volumes with MilliQ water. The lids were parafilmed to minimise evaporation. The reactions were incubated on the Thermomixer C with shaking at 600 rpm at 90 °C for 2 minutes and cooled slowly to 25 °C at a rate of -1 °C/minute.

### Extension and Ligation Reactions to Resynthesise the Plasmid

The total reaction volumes varied and depended on the amount of cssDNA used. For example, when 10 µg cssDNA template is used, the total reaction volume would be 874 µL. To the annealing reactions containing the cssDNA template (final concentration = 11.44 ng/µL), each of the dNTP (final concentration = 0.8 mM), DTT (final concentration = 5 mM), T7 DNAP (final concentration = 0.05 U/µL), T4 DNA Ligase (final concentration = 0.05 Weiss units/µL) were added. The reactions were made up to the desired volumes with MilliQ water. The reactions were incubated on the Thermomixer C with shaking at 600 rpm at 20 °C for 3 hours and kept at 4 °C overnight.

### Assessing Dinucleotides at the Ligation Junction

Each primer (0.23 pmol, **Supplementary Table 1**) was phosphorylated similarly to above except a final concentration of 0.6 U/µL was used for the T4 PNK in a total reaction volume of 5 µL. The annealing reaction was performed similarly to above except 67 ng of cssDNA template was annealed to .23 pmol phosphorylated primer in a 62.5 µL total reaction volume. The extension and ligation reactions were performed similarly to above for each annealed mix, except a 75 µL total reaction volume was used. The reactions were then lyophilised to dryness, reconstituted in MilliQ water (20 µL) and analysed on an agarose gel.

### Gyrase Reaction to Confirm Ligation

The reactions were performed in 0.2 mL PCR tubes and consisted of the ligated plasmid (100 ng), Gyrase (0.5 µL), Gyrase Reaction Buffer (4.0 µL, 5X), and made up to 20 µL with MilliQ water. The reactions were incubated on the thermal cycler at 37 °C for 1 hour and heat-deactivated at 70 °C for 30 minutes.

### DpnI Reaction to Remove Unmodified Plasmids

After extension, the amount of DNA doubled. The total reaction volumes varied and depended on the amount of dsDNA used. For example, when 1 µg dsDNA template is used, the total reaction volume would be 150 µL. To the extension and ligation reaction tubes containing the ligated plasmid (final concentration = 6.7 ng/µL), magnesium chloride (final concentration = 10 mM), Tris (pH 7.4, final concentration = 50 mM), and DpnI (final concentration = 0.06 U/µL) were added. The reactions were made up to the desired volumes with MilliQ water. The reactions were incubated on the Thermomixer C with shaking at 600 rpm at 37 °C for 5 minutes. EDTA (pH 8.0, final concentration = 20 mM) was quickly added to stop the DpnI reaction. The reactions were lyophilised to dryness.

The reactions were resuspended in MilliQ water and purified using a Genejet PCR column. The plasmids were eluted twice, each time with 50 µL MilliQ water. Ethanol precipitation was performed to remove any residual contaminants. Briefly, sodium acetate (10 µL or tenth volumes, 3 M, pH 5.2) and ethanol (300 µL or 3 volumes, 99.8%) were added to the tubes. These were vortexed and kept at -80 °C for 1-2 hours or -20 °C overnight. The tubes were centrifuged at 15,000 rpm at 4 °C for 30 minutes. The supernatant was carefully removed whilst avoiding disturbing the pellets. Ethanol (1 mL, 70%) was added and the tubes were flicked and re-centrifuged at the above settings for 10 minutes. The supernatant was removed and the pellets were dried under vacuum on a speed vac for 10 minutes. Dried DNA pellets were re-suspended in Tris (volumes depended on the reaction scale, 10 mM, pH 8.0). The reactions were Nanodropped to determine purity, whilst the concentrations were estimated using Pico488 dye.

### Measuring Concentrations with Pico488 Dye

As PCB plasmids contained 2-nitrobenzyl groups, which absorbed at 260 nm, their concentrations were determined using Pico488 dye. To ensure consistency, concentrations of the native and amine plasmids were also quantified using Pico488 before cell-free reactions and transfection. The standards were prepared using a dsDNA quantitative standard (Lumiprobe). Standards and samples were prepared according to the manufacturer’s protocol. The solutions were transferred to a clear lid 96-well flat bottom with a low evaporation lid and centrifuged at 1,250 rpm for 1 minute to remove any bubbles. The fluorescence was measured on a Tecan Infinite M1000 Pro fluorescence plate reader. Plates were shaken at 654 rpm for 10 seconds and the fluorescence was measured with λ_ex_ = 503 nm, λ_em_ = 525 nm, and bandwidth = 5 nm. The readings were used to plot the calibration curve. The equation for the line of best fit was used to calculate the concentrations.

### Binding Streptavidin

The total reaction volumes depended on the mass of dsDNA used. For example, when 1 µg dsDNA template is used, the total reaction volume would be 20 µL. The molar mass of dsDNA was calculated based on the length of the plasmid and assuming the mass of each base pair was 660. The molar amount of plasmid, and hence PCB could then be calculated, based on the number of PCBs on the PCB oligo (7 for the T7 promoter and 9 for the CMV promoter). A mass of 52.8 kDa was assumed for the monovalent and wildtype tetravalent streptavidin.

Reactions were performed in Protein LoBind tubes and consisted of the PCB plasmid (final concentration = 50.0 ng/µL), either MVS or WTS (3 molar equivalents relative to the amount of PCB sites), Tris (pH 7.4, final concentration = 20 mM), and made up to the desired volumes with MilliQ water. The reactions were incubated on the Thermomixer C with shaking at 600 rpm at 26 °C overnight.

### UV Irradiation

Cell-free reactions were assembled and irradiated immediately before incubation at 37 °C. Cells were irradiated 4 hours after transfection. The lamp (365 nm, 700 mA, M365-L2, Thorlabs), equipped with a collimation adapter (COP5-A, Thorlabs) was collimated at a distance of ∼22.5 cm (from the base of the lamp to the bottom of the tube/plate). The UV intensity was measured with a photodiode power sensor (200-1100 nm, 50 mW, Ø = 9.5 mm, S120VC, Thorlabs) equipped with a power meter console (PM100A, Thorlabs). The sensor measured UV power in mW. These values were then converted into irradiance units by dividing the power by the area of the aperture in cm^2^. The zone of irradiation was large enough to irradiate up to 4 tubes/wells simultaneously. An irradiance of 2.47 (for the T7 promoter) or 9.88 mW cm^-2^ (for the CMV promoter) was applied for 10 minutes to uncage the LA-plasmids. To ensure consistency, conditions not receiving UV were kept at room temperature whilst the +UV conditions were being irradiated. Cells were re-incubated at 37 °C and 5% CO_2_ for 2 days.

### IVT Reactions of the T7 mV_nicks Plasmid

The reactions were assembled in 0.2 mL PCR tubes and consisted of the plasmid template (200 ng), RNAPol Reaction Buffer (2 µL, 10X), each of the NTP (0.1 µL, 100 mM), RNase inhibitor (0.50 µL, 40 U/µL), T7 RNAP (2 µL, 50 U/µL), and made up to 20 µL with MilliQ water. Reactions receiving UV were irradiated as above. They were then incubated on the thermal cycler at 37 °C for 1 hour. DNase I (1 µL) was then added, and the reactions were incubated on the thermal cycler at 37 °C for another 10 minutes to remove the DNA template. To stop the reactions, EDTA (1.10 µL, 100 mM) was added, and the reactions were heat-deactivated on the thermal cycler at 75 °C for 10 minutes. 15 µL of the reactions were mixed with RNA Loading Dye (15 µL, 2X) and denatured on the thermal cycler at 90 °C for 2 minutes, kept on ice for 2 minutes and analysed on an agarose gel. All gel equipment was cleaned with RNaseZap beforehand to remove RNase contaminants.

### CFPS Reactions of the T7 mV_nicks Plasmid

Reactions were performed in 0.2 mL PCR tubes and consisted of the plasmid (100 ng), NEBexpress® S30 Synthesis Extract (2.4 µL), Protein Synthesis Buffer (5.0 µL, 2X), GamS Nuclease Inhibitor (0.2 µL, 1.5 µg/µL), Murine RNase Inhibitor (0.2 µL, 40 U/µL), T7 RNAP (0.2 µL, 450 U/µL), and made up to 10 µL with MilliQ water. Conditions requiring UV were irradiated as above. The reactions were incubated on the thermal cycler at 37 °C for 3 hours and kept at 4 °C for 30 minutes. The reactions were made up to 40 µL with MilliQ water and transferred to a 384-well black, flat-bottom optiplate. The plate was centrifuged at 1,250 rpm for 1 minute to remove any bubbles. The fluorescence was measured on a Tecan Infinite M1000 Pro fluorescence plate reader. Plates were shaken at 654 rpm for 10 seconds and the fluorescence was measured with λ_ex_ = 515 nm, λ_em_ = 527 nm, and bandwidth = 5 nm. Data were plotted on Origin.

### Maintaining HEK293T Cells

The full media contained Fluorobrite DMEM, FBS (10 % v/v), Non-Essential Amino Acids (1X), and GlutaMAX (1X). Cell stocks were kept in liquid nitrogen. Cells were maintained for 20-25 passages before a new stock was thawed and immediately revived. Cells were passaged every 3 to 4 days, or earlier if confluency exceeded 80%.

To passage, the media was removed. PBS Buffer (10 mL) was gently added and the flask was swirled and PBS was removed. Trypsin (2 mL) was added and the flask was incubated at 37 °C, 5% CO_2_ for ∼1 minute. When the majority of cells detached, trypsin was neutralised with full media (8 mL) and the solution was transferred to a 15-mL Falcon tube. The cells were centrifuged at 1,000 rpm for 3 minutes. The supernatant was carefully removed without disturbing the cell pellet. The cells were re-suspended in full media (10 mL) and split in a 1:5-1:10 ratio depending on the cell density and the user’s need. Cells were incubated at 37 °C and 5% CO_2_.

### Cell Viability Assay

The CellTiter-Glo® Reagent was prepared according to the manufacturer’s instructions and 2 mL aliquots in 15-mL Falcon tubes were wrapped in tin foil and kept at -20 °C. When needed, the reagent was thawed at room temperature and used immediately.

After cells were resuspended in full media (see above), the cell number was determined using the Countess II FL cell counter (Thermo Fisher Scientific). 10 µL of the suspension was mixed with Tryptan Blue (10 µL) and two 10-µL measurements were recorded. The average was taken and used to prepare the working solution for plating cells. The volume of cells required was transferred to a 50-mL Falcon tube and made up to the desired volume with full media to give a final concentration of 6.0 * 10^2^ cells/µL. 100 µL (6.0 * 10^4^ cells/well) was seeded onto a 96-well plate with a flat bottom and a low evaporation lid and kept at 37 °C and 5% CO_2_. After growth to >80% confluency within 24 hours, cells were irradiated with the conditions in **SI Figure 10** using the UV set-up described previously and re-incubated at 37 °C and 5% CO_2_.

The following day, the CellTiter Reagent (100 µL) was added to each well and the wells were incubated on the Thermomixer C with shaking at 600 rpm at 20 °C for 2 minutes to induce cell lysis. The plates were incubated under tin foil at room temperature for 10 minutes to stabilise the luminescence signal. Chemiluminescence was measured on a Tecan Infinite M1000 Pro fluorescence plate reader. The plates were shaken at 654 rpm for 10 seconds and the luminescence was measured using the luminescence scan mode between 559-561 nm. Values at 560 nm were used for analysis and data were plotted on Origin.

### Transfection

Before seeding cells, the wells were coated with Poly-D-Lysine (50 µL) for 5 minutes and washed twice with PBS (100 µL each time). The cells were counted and plated as above, except 5.5 * 10^4^ cells in 180 µL media were used to seed each well. The cells were incubated at 37 °C and 5% CO_2_ overnight.

Transfection with Lipofectamine™ 3000 was performed according to the manufacturer’s protocol. A 2:1 Lipofectamine 3000:DNA ratio, and 100 ng DNA were used for each well. A Lipofectamine 3000 mastermix and separate DNA mastermixes were prepared and made up to the desired volumes with MilliQ water. The Lipofectamine 3000 mastermix was added to each DNA mastermix and vortexed. The reactions were incubated at room temperature for 15 minutes to form lipoplexes. 12.40 µL of the reaction was transfected into each well and the solution was homogenised by gently rocking the plates in a circular motion. The plates were incubated at 37 °C and 5% CO_2_ for 4 hours, UV was then applied as above, and the plates were re-incubated for 2 days.

### RNA extraction

A working solution of Buffer RLT containing beta-mercaptoethanol (βME) was prepared fresh (10 µL βME per mL Buffer RLT).

The media in each well was removed and the wells were washed with PBS (100 µL). Trypsin (100 µL) was added to each well and the plates were kept at 37 °C and 5% CO_2_ until the majority of cells detached. Trypsin was neutralised with full media (100 µL) and the solutions were transferred to 1.5 mL Eppendorf tubes. RNA extraction was performed using RNeasy Mini kit according to the manufacturer’s protocol. RNA was eluted with RNase-free water (50 µL). RNA was ethanol-precipitated as above and resuspended in RNase-free water (20 µL). Concentrations were measured using the NanoDrop and the 260/230 ratios were typically >1.90. RNA was stored at -80 °C.

### ezDNase and cDNA Synthesis

gDNA was removed from RNA preparations with ezDNase according to the manufacturer’s protocol except the reaction was scaled to 6 µL total volume and 60 ng RNA was used.

To each tube, Superscript IV Vilo MM (2.40 µL, 5X) was added and the reactions were made up to 12.0 µL with MilliQ water. The reactions were mixed and incubated on the thermal cycler with the following program: annealed at 25 °C for 10 minutes, reverse transcribed at 50 °C for 10 minutes, and heat-deactivated at 85 °C for 5 minutes. The cDNA was stored at -80 °C.

### RT-qPCR

Reactions were performed in 0.2 mL PCR tubes and consisted of the cDNA (2 µL), mNG_qPCR_FOR (0.4 µL, 10 µM), mNG_qPCR_REV (0.4 µL, 10 µM, **Supplementary Table 3**), Fast SYBR Green MM (10 µL, 2X), and made up to 20 µL with MilliQ water to amplify mNG. Normalisation with GAPDH was performed analogously using GAPDH_FOR_Origene and GAPDH_REV_IDT as primers (**Supplementary Table 3**). Reactions were cycled on the CFX96 real-Time System (C1000 Touch™ Thermalcycler, Bio-Rad) with SYBR Green monitoring using the following parameters: denaturation at 95 °C for 20 seconds, 30 cycles [95 °C for 3 seconds, 60 °C for 30 seconds]. The ΔΔCt method was used to quantify the transcription fold change. The data were set relative to ‘amine plasmid -UV’ condition and plotted using Origin.

### Plate Reader of CMV mNG Plasmids

After 2 days, the fluorescence was measured on the plate reader. Plates were shaken at 654 rpm for 10 seconds and the fluorescence was measured with λ_ex_ = 504 nm, λ_em_ = 517 nm, and bandwidth = 5 nm. Data were background-subtracted and plotted on Origin.

### Epifluorescence Microscopy of CMV mNG Plasmids

Cells were imaged using a Leica DMi8 inverted epifluorescence microscope with a 10x objective lens. Cells were imaged with the brightfield and GFP channels. Analysis was performed with ImageJ.

## Acknowledgements

We would like to thank Prof. M. Howarth and Dr. M. Fairhead for kindly providing the monovalent streptavidin, and Professor Dame Carol Robison and Dr. Haiping Tang for providing the HEK293T cell line.

K.C. is supported through the Synthetic Biology Centre for Doctoral Training – EPSRC (EP/L016494/1) and CRUK funding (C2195/A27450).

M. J. B. is supported by a Royal Society University Research Fellowship and an EPSRC New Investigator Award (EP/V030434/1).

## Contributions

K.C. and M.J.B. designed the project. K.C. designed, performed, and analysed the experiments, with contributions from M.J.B. K.C. and M.J.B. wrote the paper.

