## Supplementary Information for "Sequence-independent, site-specific incorporation of chemical modifications to generate light-activated plasmids"

### DNA Sequences

**Supplementary Table 1. Primers used to assess dinucleotides at the ligation junction**

The primers followed the format X_Y_FOR or X_Y_phos where X represents the nucleotide at the 3'-end and Y represents the nucleotide at the 5'-end of the ligation junction. 'phos' primers were already 5' phosphorylated.

| Name | Sequence |
|---|---|
| C_C_FOR | CTGAAGCTCATCTGCACC |
| G_T_FOR | TCCAGGAGCGCACCATCTTC |
| T_C_FOR | CAAGGACGACGGCAACTAC |
| G_C_FOR | CATCGAGCTGAAGGGCATC |
| C_T_FOR | TTCAAGGAGGACGGCAACATCC |
| G_G_FOR | GGAATTCTCGAGTAAGGTTAACC |
| T_A_FOR | ATATCCGGAAGCTTGGCACTGG |
| G_A_FOR | AGCGGTATCAGCTCACTC |
| C_A_FOR | AAGTCAGAGGTGGCGAAACCC |
| A_A_FOR | AGCGTGGCGCTTTCTCATAGC |
| T_G_phos | GTAGGTCGTTCGCTCCAAGC |
| A_G_FOR | GTAAACTTGGTCTGACAGTTACC |
| A_T_FOR | TCTCAGCGATCTGTCTATTTCG |
| T_T_FOR | TCATCCATAGTTGCCTGAC |
| C_G_phos | GAGTTGCTCTTGCCCGGCGTC |
| A_C_phos | CACGGAAATGTTGAATACTC |

**Supplementary Table 2. mV_nicks and mNG_BbvCI plasmid sequences**

| Plasmid | Sequence |
|---|---|
| mV_nicks | GCTAGTGGTGCTAGCCCCGCGAAATTAATACGACTCACTATAGG<br>GTCTAGAAATAATTTTGTTTAACTTTAAGAAGGAGGTATACATA<br>TGGTGAGCAAGGGCGAGGAGCTGTTCACCGGGGTGGTGCCCAT<br>CCTGGTCGAGCTGGACGGCGACGTAAACGGCCACAAGTTCAGC<br>GTGTCCGGCGAGGGCGAGGGCGATGCCACCTACGGCAAGCTGA<br>CCCTGAAGCTCATCTGCACCACCGGCAAGCTGCCCGTGCCCTGG<br>CCCACCCTCGTGACCACCCTCGGCTACGGCCTGCAGTGCTTCGC<br>CCGCTACCCCGACCACATGAAGCAGCACGACTTCTTCAAGTCCG<br>CCATGCCCGAAGGCTACGTCCAGGAGCGCACCATCTTCTTCAAG<br>GACGACGGCAACTACAAGACCCGCGCCGAGGTGAAGTTCGAGG<br>GCGACACCCTGGTGAACCGCATCGAGCTGAAGGGCATCGACTT<br>CAAGGAGGACGGCAACATCCTGGGGCACAAGCTGGAGTACAAC<br>TACAACAGCCACAACGTCTATATCACCGCCGACAAGCAGAAGA<br>ACGGCATCAAGGCCAACTTCAAGATCCGCCACAACATCGAGGA<br>CGGCGGCGTGCAGCTCGCCGACCACTACCAGCAGAACACCCCC<br>ATCGGCGACGGCCCCGTGCTGCTGCCCGACAACCACTACCTGAG<br>CTACCAGTCCAAGCTGAGCAAAGACCCCAACGAGAAGCGCGAT<br>CACATGGTCCTGCTGGAGTTCGTGACCGCCGCCGGGATCACTCT<br>CGGCATGGACGAGCTGTACAAGTAATGACCTCAGCGGATCCGC<br>TCTTCCGGGAATTCTCGAGTAAGGTTAACCTGCAGGAGGCCTTT<br>AATTAAGGTGGTGCGGCCGCGCTAGCGGTCCCGGGGGATCGAT<br>CCGGCTGCTAACAAAGCCCGAAAGGAAGCTGAGTTGGCTGCTG<br>CCACCGCTGAGCAATAACTAGCATAACCCCTTGGGGCCTCTAAA |

| | |
|---|---|
| | CGGGTCTTGAGGGGTTTTTGCTGAAAGGAGGAACTATATCCGG<br>AAGCTTGGCACTGGCCGACCGGGGTCGAGCACTGACTCGCTGC<br>GCTCGGTCGTTCGGCTGCGGCGAGCGGTATCAGCTCACTCAAAG<br>GCGGTAATACGGTTATCCACAGAATCAGGGGATAACGCAGGAA<br>AGAACATGTGAGCAAAAGGCCAGCAAAAGGCCAGGAACCGTA<br>AAAAGGCCGCGTTGCTGGCGTTTTTCCATAGGCTCCGCCCCCCT<br>GACGAGCATCACAAAAATCGACGCTCAAGTCAGAGGTGGCGAA<br>ACCCGACAGGACTATAAAGATACCAGGCGTTTCCCCCTGGAAG<br>CTCCCTCGTGCGCTCTCCTGTTCCGACCCTGCCGCTTACCGGATA<br>CCTGTCCGCCTTTCTCCCTTCGGGAAGCGTGGCGCTTTCTCATAG<br>CTCACGCTGTAGGTATCTCAGTTCGGTGTAGGTCGTTCGCTCCA<br>AGCTGGGCTGTGTGCACGAACCCCCCGTTCAGCCCGACCGCTGC<br>GCCTTATCCGGTAACTATCGTCTTGAGTCCAACCCGCTAAGACA<br>CGACTTATCGCCACTGGCAGCAGCCACTGGTAACAGGATTAGCA<br>GAGCGAGGTATGTAGGCGGTGCTACAGAGTTCTTGAAGTGGTG<br>GCCTAACTACGGCTACACTAGAAGAACAGTATTTGGTATCTGCG<br>CTCTGCTGAAGCCAGTTACCTTCGGAAAAAGAGTTGGTAGCTCT<br>TGATCCGGCAAACAAACCACCGCTGGTAGCGGTGGTTTTTTTGT<br>TTGCAAGCAGCAGATTACGCGCAGAAAAAAAGGATCTCAAGAA<br>GATCCTTTGATCTTTTCTACGGGGTCTGACGCTCAGTGGAACGA<br>AAACTCACAGATCCGGGATTTTGGTCATGAGATTATCAAAAAGG<br>ATCTTCACCTAGATCCTTTTAAATTAAAAATGAAGTTTTAAATC<br>AATCTAAAGTATATATGAGTAAACTTGGTCTGACAGTTACCAAT<br>GCTTAATCAGTGAGGCACCTATCTCAGCGATCTGTCTATTTCGTT<br>CATCCATAGTTGCCTGACTCCCCGTCGTGTAGATAACTACGATA<br>CGGGAGGGCTTACCATCTGGCCCCAGTGCTGCAATGATACCGCG<br>AGACCCACGCTCACCGGCTCCAGATTTATCAGCAATAAACCAGC<br>CAGCCGGAAGGGCCGAGCGCAGAAGTGGTCCTGCAACTTTATC<br>CGCCTCCATCCAGTCTATTAATTGTTGCCGGGAAGCTAGAGTAA<br>GTAGTTCGCCAGTTAATAGTTTGCGCAACGTTGTTGCCATTGCT<br>ACAGGCATCGTGGTGTCACGCTCGTCGTTTGGTATGGCTTCATT<br>CAGCTCCGGTTCCCAACGATCAAGGCGAGTTACATGATCCCCCA<br>TGTTGTGCAAAAAAGCGGTTAGCTCCTTCGGTCCTCCGATCGTT<br>GTCAGAAGTAAGTTGGCCGCAGTGTTATCACTCATGGTTATGGC<br>AGCACTGCATAATTCTCTTACTGTCATGCCATCCGTAAGATGCTT<br>TTCTGTGACTGGTGAGTACTCAACCAAGTCATTCTGAGAATAGT<br>GTATGCGGCGACCGAGTTGCTCTTGCCCGGCGTCAATACGGGAT<br>AATACCGCGCCACATAGCAGAACTTTAAAAGTGCTCATCATTGG<br>AAAACGTTCTTCGGGGCGAAAACTCTCAAGGATCTTACCGCTGT<br>TGAGATCCAGTTCGATGTAACCCACTCGTGCACCCAACTGATCT<br>TCAGCATCTTTTACTTTCACCAGCGTTTCTGGGTGAGCAAAAAC<br>AGGAAGGCAAAATGCCGCAAAAAAGGGAATAAGGGCGACACG<br>GAAATGTTGAATACTCATACTCTTCCTTTTTCAATATTATTGAAG<br>CATTTATCAGGGTTATTGTCTCATGAGCGGATACATATTTGAAT<br>GTATTTAGAAAAATAAACAAATAGGGGTTCCGCGCACATTTCCC<br>CGAAAAGT |
| mNG_BbvCI | TCGACCGCCAATTCAATATGGCGTATATGGACTCATGCCAATTC<br>AATATGGTGGATCTGGACCTGTGCCAATTCAATATGGCGTATAT<br>GGACTCGTGCCAATTCAATATGGTGGATCTGGACCCCAGCCAAT<br>TCAATATGGCGGACTTGGCACCATGCCAATTCAATATGGCGGAC |

| | |
|---|---|
| | CTGGCACTGTGCCAACTGGGGAGGGGTCTACTTGGCACGGTGCC<br>AAGTTTGAGGAGGGGTCTTGGCCCTGTGCCAAGTCCGCCATATT<br>GAATTGGCATGGTGCCAATAATGGCGGCCATATTGGCTATATGC<br>CAGGATCAATATATAGGCAATATCCAATATGGCCCTATGCCAAT<br>ATGGCTATTGGCCAGGTTCAATACTATGTATTGGCCCTATGCCA<br>TATAGTATTCCATATATGGGTTTTCCTATTGACGTAGATAGCCCC<br>TCCCAATGGGCGGTCCCATATACCATATATGGGGCTTCCTAATA<br>CCGCCCATAGCCACTCCCCCATTGACGTCAATGGTCTCTATATA<br>TGGTCTTTCCTATTGACGTCATATGGGCGGTCCTATTGACGTATA<br>TGGCGCCTCCCCCATTGACGTCAATTACGGTAAATGGCCCGCCT<br>GGCTCAATGCCCATTGACGTCAATAGGACCACCCACCATTGACG<br>TCAATGGGATGGCTCATTGCCCATTCATATCCGTTCTCACGCCCC<br>CTATTGACGTCAATGACGGTAAATGGCCCACTTGGCAGTACATC<br>AATATCTATTAATAGTAACTTGGCAAGTACATTACTATTGGAAG<br>TACGCCAGGGTACATTGGCAGTACTCCCATTGACGTCAATGGCG<br>GTAAATGGCCCGCGATGGCTGCCAAGTACATCCCCATTGACGTC<br>AATGGGGAGGGGCAATGACGCAAATGGGCGTTCCATTGACGTA<br>AATGGGCGGTAGGCGTGCCTAATGGGAGGTCTATATAAGCAAT<br>GCTCGTTTAGGGAACCGCCATTCTGCCTGGGGACGTCGGAGCAA<br>GCTTGATTTAGGTGACACTATAGACTTGTTCTTTTTGCAGGATCT<br>ACCATGGTGAGCAAGGGCGAGGAGGATAACATGGCCTCTCTCC<br>CAGCGACACATGAGTTACACATCTTTGGCTCCATCAACGGTGTG<br>GACTTTGACATGGTGGGTCAGGGCACCGGCAATCCAAATGATG<br>GTTATGAGGAGTTAAACCTGAAGTCCACCAAGGGTGACCTCCA<br>GTTCTCCCCCTGGATTCTGGTCCCTCATATCGGGTATGGCTTCCA<br>TCAGTACCTGCCCTACCCTGACGGGATGTCGCCTTTCCAGGCCG<br>CCATGGTAGATGGCTCCGGATACCAAGTCCATCGCACAATGCAG<br>TTTGAAGATGGTGCCTCCCTTACTGTTAACTACCGCTACACCTAC<br>GAGGGAAGCCACATCAAAGGAGAGGCCCAGGTGAAGGGGACT<br>GGTTTCCCTGCTGACGGTCCTGTGATGACCAACTCGCTGACCGC<br>TGCGGACTGGTGCAGGTCGAAGAAGACTTACCCCAACGACAAA<br>ACCATCATCAGTACCTTTAAGTGGAGTTACACCACTGGAAATGG<br>CAAGCGCTACCGGAGCACTGCGCGGACCACCTACACCTTTGCCA<br>AGCCAATGGCGGCTAACTATCTGAAGAACCAGCCGATGTACGT<br>GTTCCGTAAGACGGAGCTCAAGCACTCCAAGACCGAGCTCAAC<br>TTCAAGGAGTGGCAAAAGGCCTTTACCGATGTGATGGGCATGG<br>ACGAGCTGTACAAGGGATCTGGATCCCATCGATTCGAATTCAAG<br>GCCTCTCGAGCCTCTAGAACTATAGTGAGTCGTATTACGTAGAT<br>CCAGACATGATAAGATACATTGATGAGTTTGGACAAACCACAA<br>CTAGAATGCAGTGAAAAAAATGCTTTATTTGTGAAATTTGTGAT<br>GCTATTGCTTTATTTGTAACCATTATAAGCTGCAATAAACAAGT<br>TAACAACAACAATTGCATTCATTTTATGTTTCAGGTTCAGGGGG<br>AGGTGTGGGAGGTTTTTTAATTCGCGGCCGCGGCGCCAATGCAT<br>TGGGCCCGGTACCCAGCTTTTGTTCCCTTTAGTGAGGGTTAATTG<br>CGCGCTTGGCGTAATCCCTCAGCATGGTCATAGCTGTTTCCTGT<br>GTGAAATTGTTATCCGCTCACAATTCCACACAACATACGAGCCG<br>GGAGCATAAAGTGTAAAGCCTGGGGTGCCTAATGAGTGAGCTA<br>ACTCACATTAATTGCGTTGCGCTCACTGCCCGCTTTCCAGTCGG<br>GAAACCTGTCGTGCCAGCTGCATTAATGAATCGGCCAACGCGCG<br>GGGAGAGGCGGTTTGCGTATTGGGCGCTCTTCCGCTTCCTCGCT |

| | |
|---|---|
| | CACTGACTCGCTGCGCTCGGTCGTTCGGCTGCGGCGAGCGGTAT<br>CAGCTCACTCAAAGGCGGTAATACGGTTATCCACAGAATCAGG<br>GGATAACGCAGGAAAGAACATGTGAGCAAAAGGCCAGCAAAA<br>GGCCAGGAACCGTAAAAAGGCCGCGTTGCTGGCGTTTTTCCATA<br>GGCTCCGCCCCCCTGACGAGCATCACAAAAATCGACGCTCAAGT<br>CAGAGGTGGCGAAACCCGACAGGACTATAAAGATACCAGGCGT<br>TTCCCCCTGGAAGCTCCCTCGTGCGCTCTCCTGTTCCGACCCTGC<br>CGCTTACCGGATACCTGTCCGCCTTTCTCCCTTCGGGAAGCGTG<br>GCGCTTTCTCATAGCTCACGCTGTAGGTATCTCAGTTCGGTGTA<br>GGTCGTTCGCTCCAAGCTGGGCTGTGTGCACGAACCCCCCGTTC<br>AGCCCGACCGCTGCGCCTTATCCGGTAACTATCGTCTTGAGTCC<br>AACCCGGTAAGACACGACTTATCGCCACTGGCAGCAGCCACTG<br>GTAACAGGATTAGCAGAGCGAGGTATGTAGGCGGTGCTACAGA<br>GTTCTTGAAGTGGTGGCCTAACTACGGCTACACTAGAAGAACAG<br>TATTTGGTATCTGCGCTCTGCTGAAGCCAGTTACCTTCGGAAAA<br>AGAGTTGGTAGCTCTTGATCCGGCAAACAAACCACCGCTGGTAG<br>CGGTGGTTTTTTTGTTTGCAAGCAGCAGATTACGCGCAGAAAAA<br>AAGGATCTCAAGAAGATCCTTTGATCTTTTCTACGGGGTCTGAC<br>GCTCAGTGGAACGAAAACTCACGTTAAGGGATTTTGGTCATGAG<br>ATTATCAAAAAGGATCTTCACCTAGATCCTTTTAAATTAAAAAT<br>GAAGTTTTAAATCAATCTAAAGTATATATGAGTAAACTTGGTCT<br>GACAGTTACCAATGCTTAATCAGTGAGGCACCTATCTCAGCGAT<br>CTGTCTATTTCGTTCATCCATAGTTGCCTGACTCCCCGTCGTGTA<br>GATAACTACGATACGGGAGGGCTTACCATCTGGCCCCAGTGCTG<br>CAATGATACCGCGAGACCCACGCTCACCGGCTCCAGATTTATCA<br>GCAATAAACCAGCCAGCCGGAAGGGCCGAGCGCAGAAGTGGTC<br>CTGCAACTTTATCCGCCTCCATCCAGTCTATTAATTGTTGCCGGG<br>AAGCTAGAGTAAGTAGTTCGCCAGTTAATAGTTTGCGCAACGTT<br>GTTGCCATTGCTACAGGCATCGTGGTGTCACGCTCGTCGTTTGG<br>TATGGCTTCATTCAGCTCCGGTTCCCAACGATCAAGGCGAGTTA<br>CATGATCCCCCATGTTGTGCAAAAAAGCGGTTAGCTCCTTCGGT<br>CCTCCGATCGTTGTCAGAAGTAAGTTGGCCGCAGTGTTATCACT<br>CATGGTTATGGCAGCACTGCATAATTCTCTTACTGTCATGCCATC<br>CGTAAGATGCTTTTCTGTGACTGGTGAGTACTCAACCAAGTCAT<br>TCTGAGAATAGTGTATGCGGCGACCGAGTTGCTCTTGCCCGGCG<br>TCAATACGGGATAATACCGCGCCACATAGCAGAACTTTAAAAG<br>TGCTCATCATTGGAAAACGTTCTTCGGGGCGAAAACTCTCAAGG<br>ATCTTACCGCTGTTGAGATCCAGTTCGATGTAACCCACTCGTGC<br>ACCCAACTGATCTTCAGCATCTTTTACTTTCACCAGCGTTTCTGG<br>GTGAGCAAAAACAGGAAGGCAAAATGCCGCAAAAAAGGGAAT<br>AAGGGCGACACGGAAATGTTGAATACTCATACTCTTCCTTTTTC<br>AATATTATTGAAGCATTTATCAGGGTTATTGTCTCATGAGCGGA<br>TACATATTTGAATGTATTTAGAAAAATAAACAAATAGGGGTTCC<br>GCGCACATTTCCCCGAAAAGTGCCACCTAAATTGTAAGCGTTAA<br>TATTTTGTTAAAATTCGCGTTAAATTTTTGTTAAATCAGCTCATT<br>TTTTAACCAATAGGCCGAAATCGGCAAAATCCCTTATAAATCAA<br>AAGAATAGACCGAGATAGGGTTGAGTGTTGTTCCAGTTTGGAAC<br>AAGAGTCCACTATTAAAGAACGTGGACTCCAACGTCAAAGGGC<br>GAAAAACCGTCTATCAGGGCGATGGCCCACTACGTGAACCATC<br>ACCCTAATCAAGTTTTTTGGGGTCGAGGTGCCGTAAAGCACTAA |

| | |
|---|---|
| | ATCGGAACCCTAAAGGGAGCCCCCGATTTAGAGCTTGACGGGGAAAGCCGGCGAACGTGGCGAGAAAGGAAGGGAAGAAAGCGAAAGGAGCGGGCGCTAGGGCGCTGGCAAGTGTAGCGGTCACGCTGCGCGTAACCACCACACCCGCCGCGCTTAATGCGCCGCTACAGGGCGCGTCCCATTCGCCATTCAGGCTGCGCAACTGTTGGGAAGGGCGATCGGTGCGGGCCTCTTCGCTATTACGCCAG |

**Supplementary Table 3. Primers used for PCR, RT-qPCR and NEEL**

Underlined sequences are the T7 promoter and TATA box. Bolded sequences are the upstream and downstream BRE and Inr elements. X represents the amino-C6-dT modifications.

| Name | Sequence |
|---|---|
| mV nick T7T FRW | TACAAGTAATGACCTCAGCGGATCCGCTCTTCCGGGAATTCTCGAGTAAGG |
| DHFR REV T7S GFP | ACAGCTCCTCGCCCTTGCTCACCATATGTATACCTCCTTCTTAAAGTTAAAC |
| GFP FRW T7S | CGGCATGGACGAGCTGTACAAGTAATGAGGATCCCGGGAATTCTCGAGTA |
| mV nick T7T REV | AATTCCCGGAAGAGCGGATCCGCTGAGGTCATTACTTGTACAGCTCG |
| mNG_BbvCI_FOR1 | GCTTGGCGTAATCCCTCAGCATGGTCATAGCTG |
| mNG_BbvCI_REV2 | CAGCTATGACCATGCTGAGGGATTACGCCAAGC |
| mNG_nick_mutagenesis_REV1 | CGAGAGGCCTTGAATTCGAATCGATG |
| mNG_nick_mutagenesis_FOR2 | CATCGATTCGAATTCAAGGCCTCTCG |
| mNG_qPCR_FOR | CGTGTTCCGTAAGACGGAGC |
| mNG_RT-qPCR_REV | CCTTGTACAGCTCGTCCATGC |
| GAPDH_FOR_Origene | GTCTCCTCTGACTTCAACAGCG |
| GAPDH_REV_IDT | TCCACCACCCTGTTGCTGTA |
| T7_eLong | GAAATTAATACGACTCACTATAGGGTCTAG |
| T7_7amines | GAAATXAAXACGACXCACXAXAGGGXCXAG |
| mNG_9amines_46b | **GGXC**XAXAXAAG**CAAXGCX**CGTTTAGGGAACCG**CCATXC** **X**GCCXGG |

**Supplementary Table 4. Primers used to synthesise the backbones and inserts for homologous recombinations**

| PCR | DNA template | For primer | Rev Primer |
|---|---|---|---|
| mV_nicks_BB | mVenus_CT | mV nick T7T FRW | DHFR REV T7S GFP |
| mV_nicks_insert | mVenus_CT | GFP FRW T7S | mV nick T7T REV |
| mNG_BbvCI_BB | pCS2-mNG-C plasmid | mNG_BbvCI_FOR1 | mNG_nick_mutagenesis_REV1 |
| mNG_BbvCI_insert | pCS2-mNG-C plasmid | mNG_BbvCI_REV2 | mNG_nick_mutagenesis_FOR2 |

**Supplementary Table 5. Backbone and insert combinations for homologous recombinations**

| Plasmid | Backbone | Insert |
|---|---|---|
| mV_nicks | mV_nicks_BB | mV_nicks_insert |
| mNG_BbvCI | mNG_BbvCI_BB | mNG_BbvCI_insert |

**Supplementary Table 6. HPLC Purification of the PCB oligos**

| Promoter | Number of modifications | Collection period/ mins | Increments/ mins | Fractions pooled |
|---|---|---|---|---|
| T7 | 7 | 24.0-25.5 | 0.15 | 5-7 |
| CMV | 9 | 24.0-25.5 | 0.15 | 3-5 |

### Supplementary Data

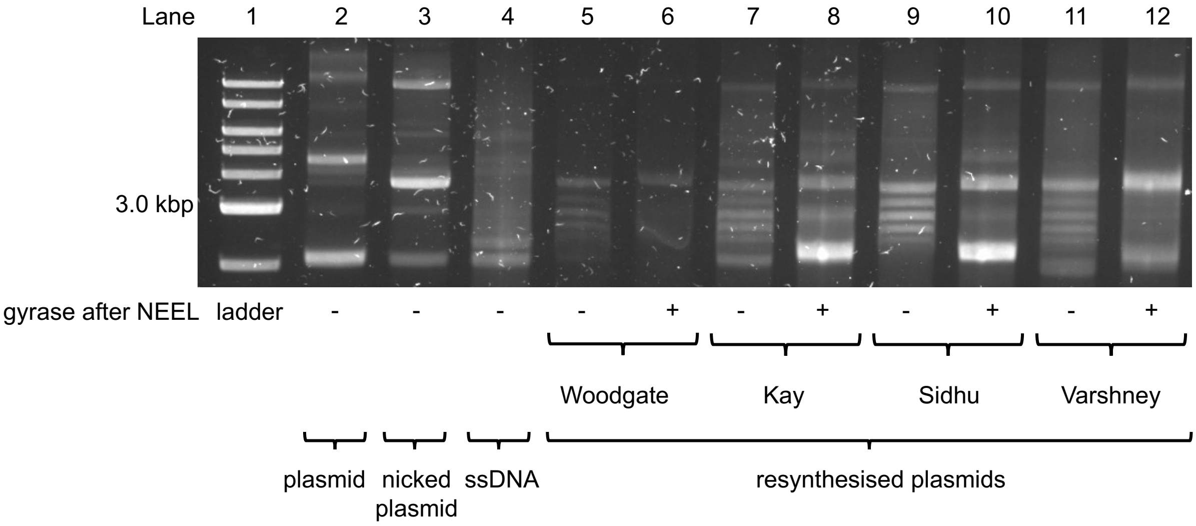

**SI Figure 1. Different Kunkel methods were employed to resynthesise the plasmid.** Kunkel mutagenesis protocols from the Woodgate,[1] Kay,[2] Sidhu,[3] and Varshney[4] laboratories were used to regenerate the plasmid. After ligation, the different plasmids were reacted with gyrase to test for ligation.

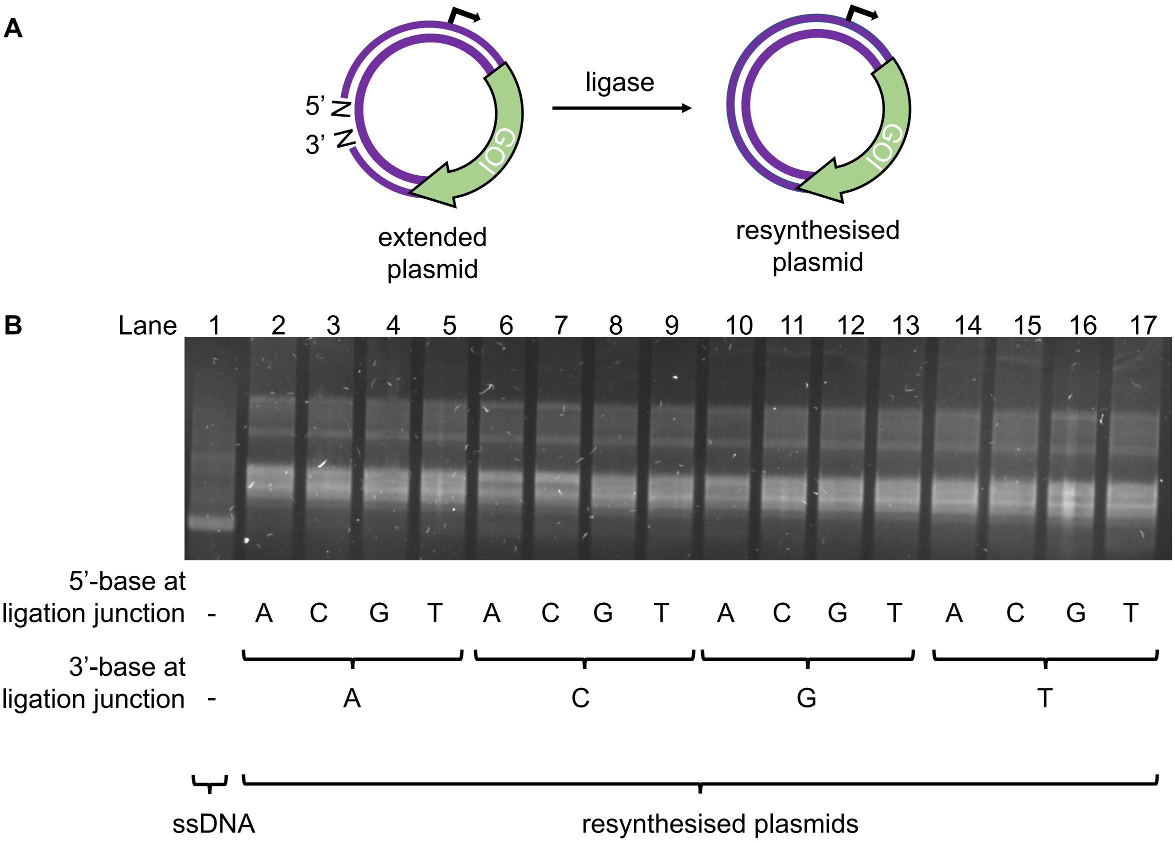

**SI Figure 2. NEEL with all possible dinucleotide combinations at the ligation junction. A.** Depiction of the reaction in (**B**). **B.** Different primers with varying $T_m$, GC content, and lengths were used for NEEL. Ligation at the different ligation junctions was equally effective.

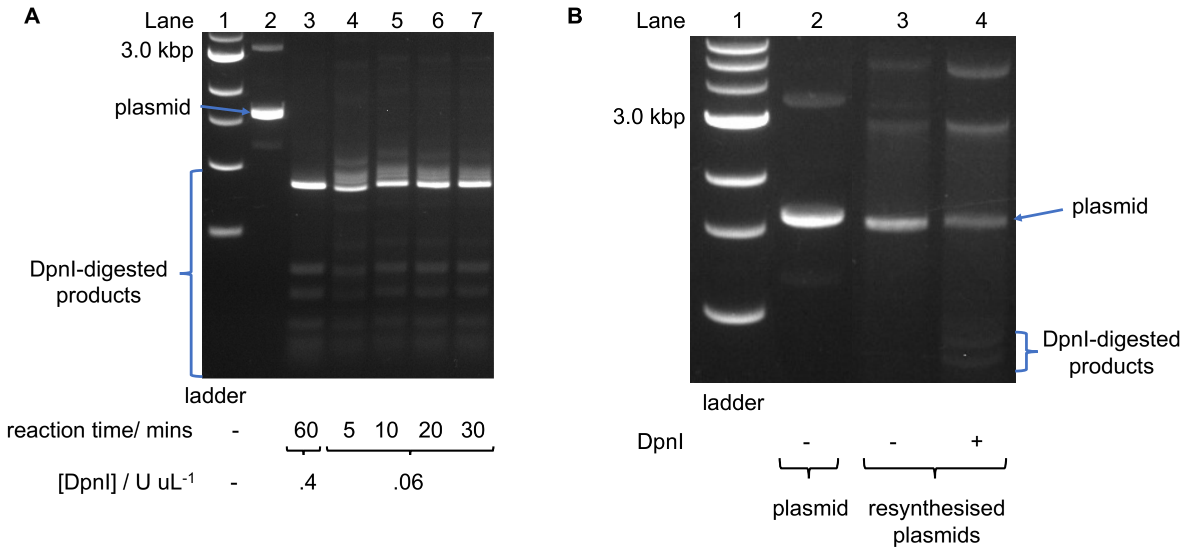

**SI Figure 3. DpnI removal of unreacted plasmids. A.** Mild DpnI conditions were optimised with native mVenus plasmid. **B.** When resynthesised plasmids were reacted with DpnI faint degraded plasmid bands appeared whilst the majority of the resynthesised plasmids were intact.

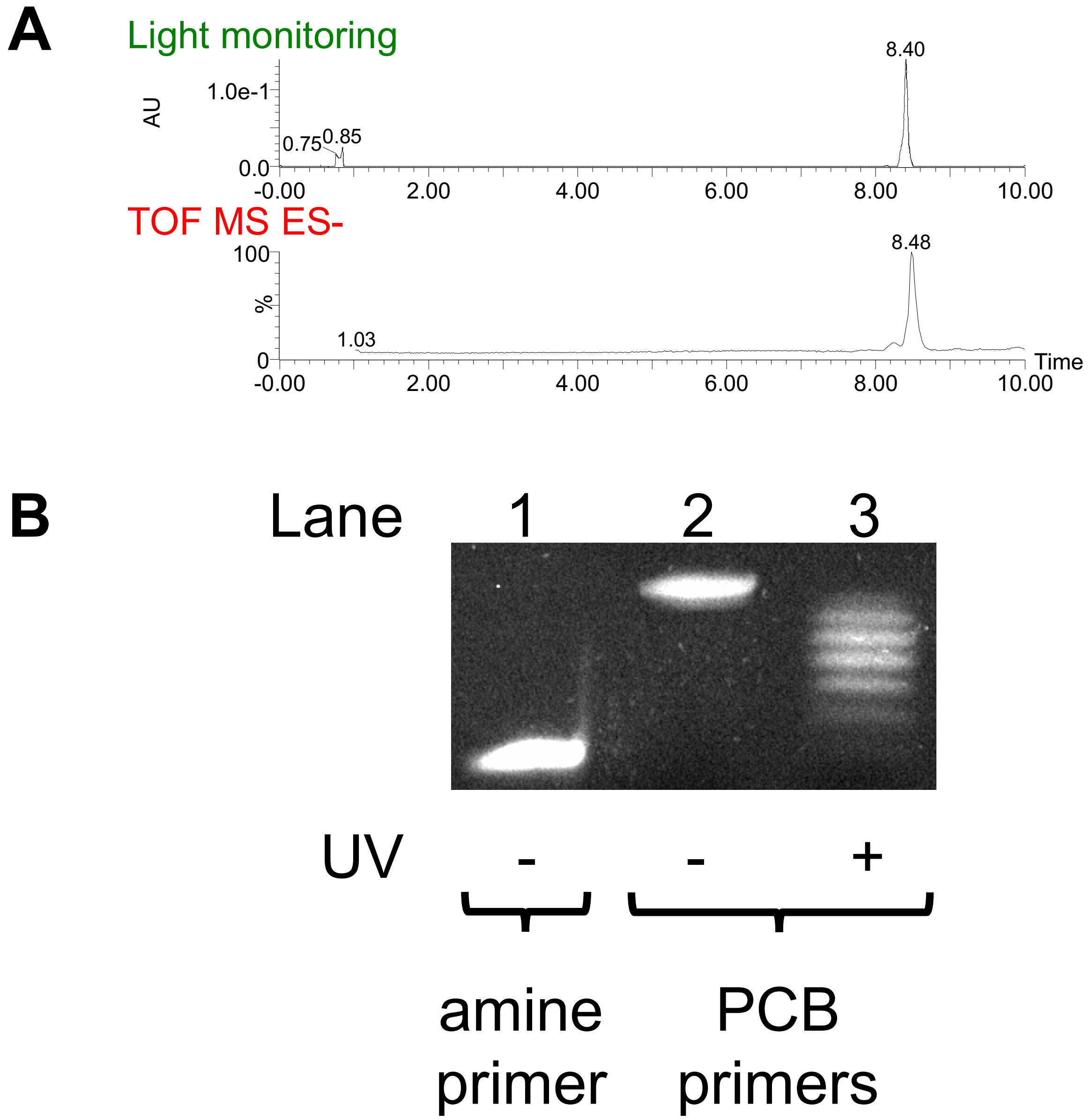

**SI Figure 4. Confirming synthesis of the T7 PCB oligo. A.** LC-MS and **B.** Denaturing PAGE were used to confirm PCB attachment. MS gave a mass of 15398, which corresponded to the 7-PCBs oligo.

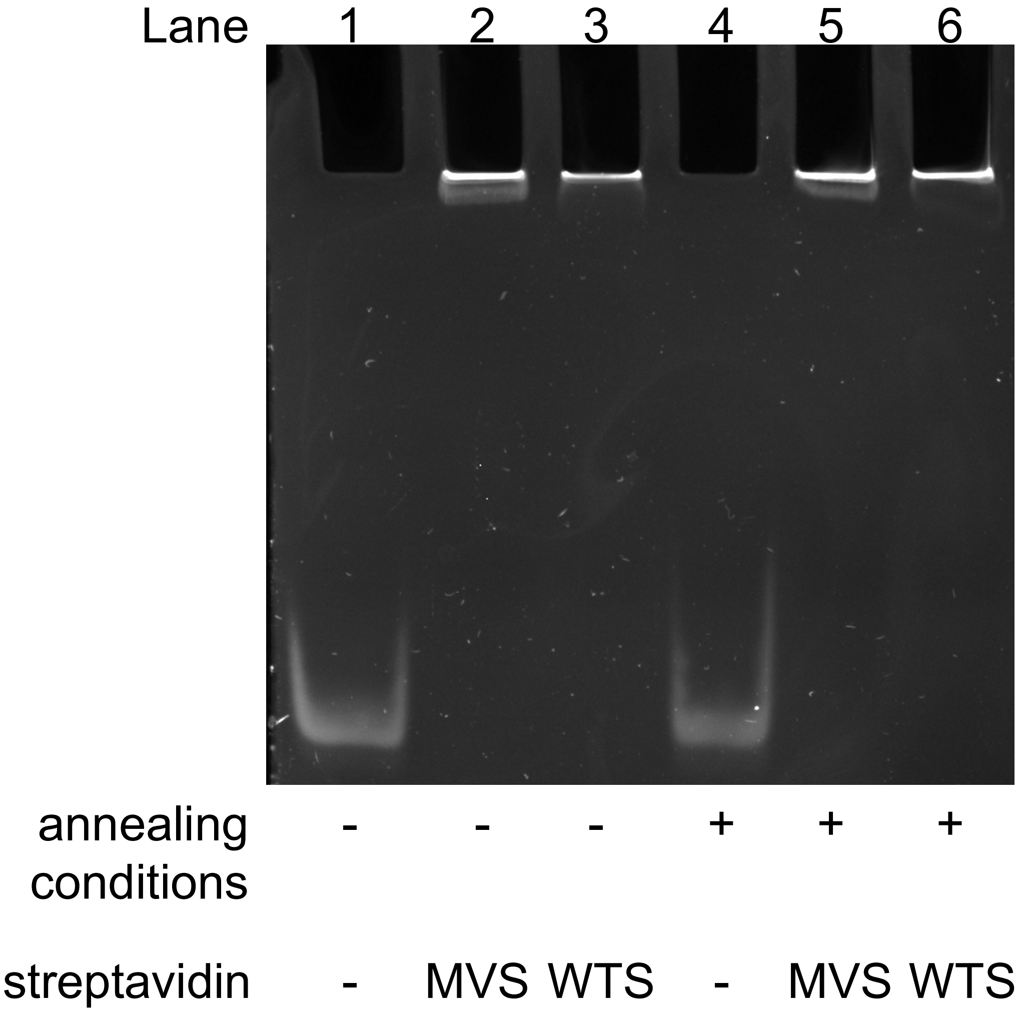

**SI Figure 5. PCB oligo was unaffected by annealing conditions.** PCB oligo was bound to either monovalent or wildtype tetravalent streptavidin (MVS and WTS, respectively) and incubated in annealing conditions.

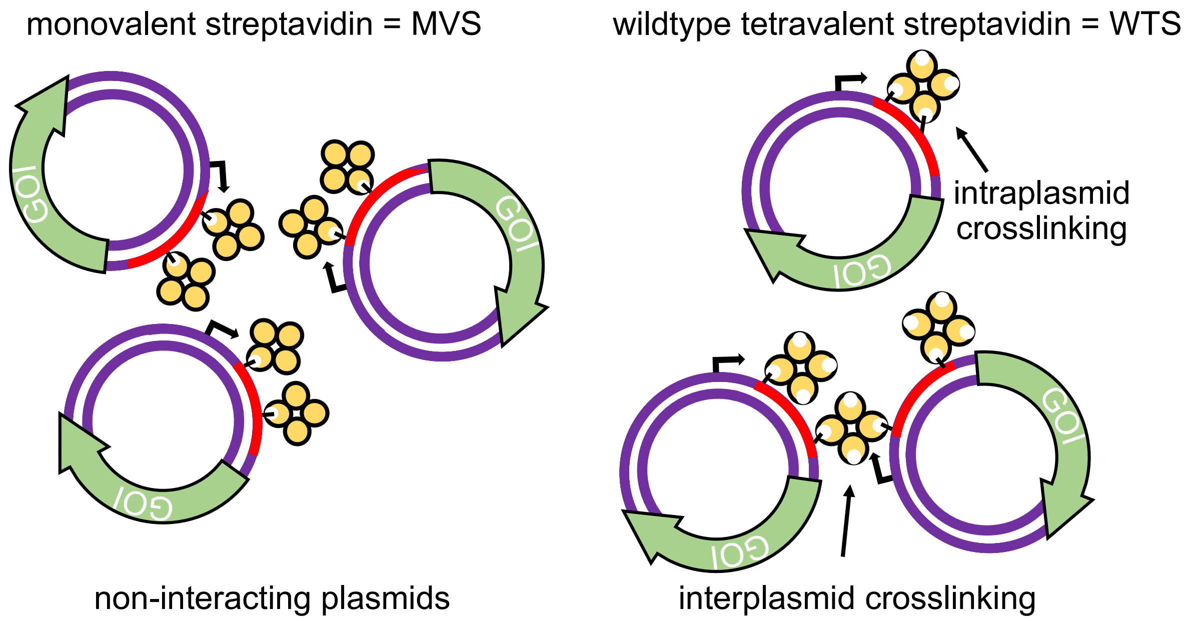

**SI Figure 6. Comparison between MVS and WTS LA-plasmids.** With only a single active subunit, MVS LA-plasmids do not interact. WTS has four active subunits, which may give rise to intra- and interplasmid crosslinking.

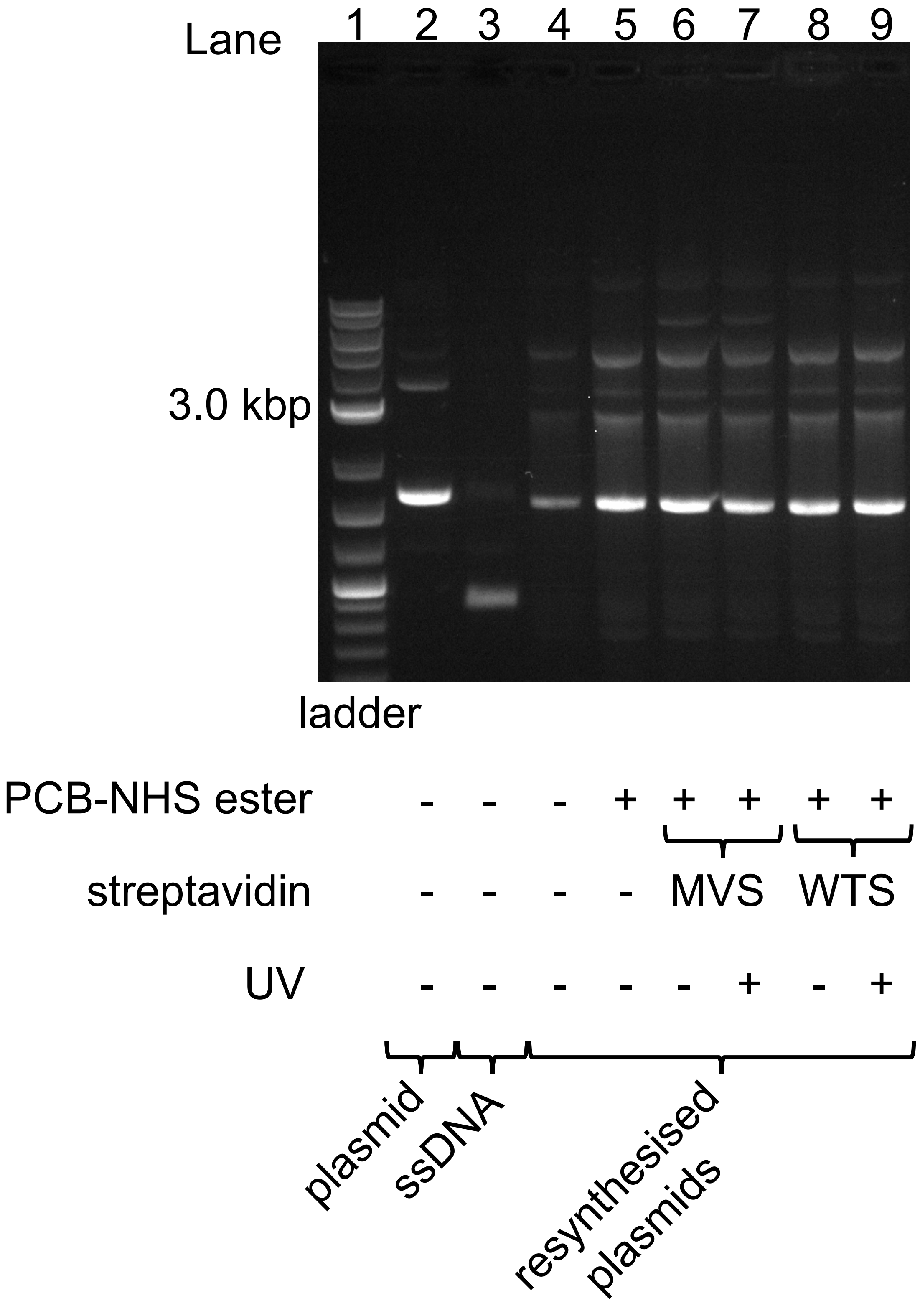

**SI Figure 7. Negative controls without amino modifications.** Plasmids were resynthesised using a regular primer without amino modifications. The absence of amines prevented PCB conjugation and thereby streptavidin binding.

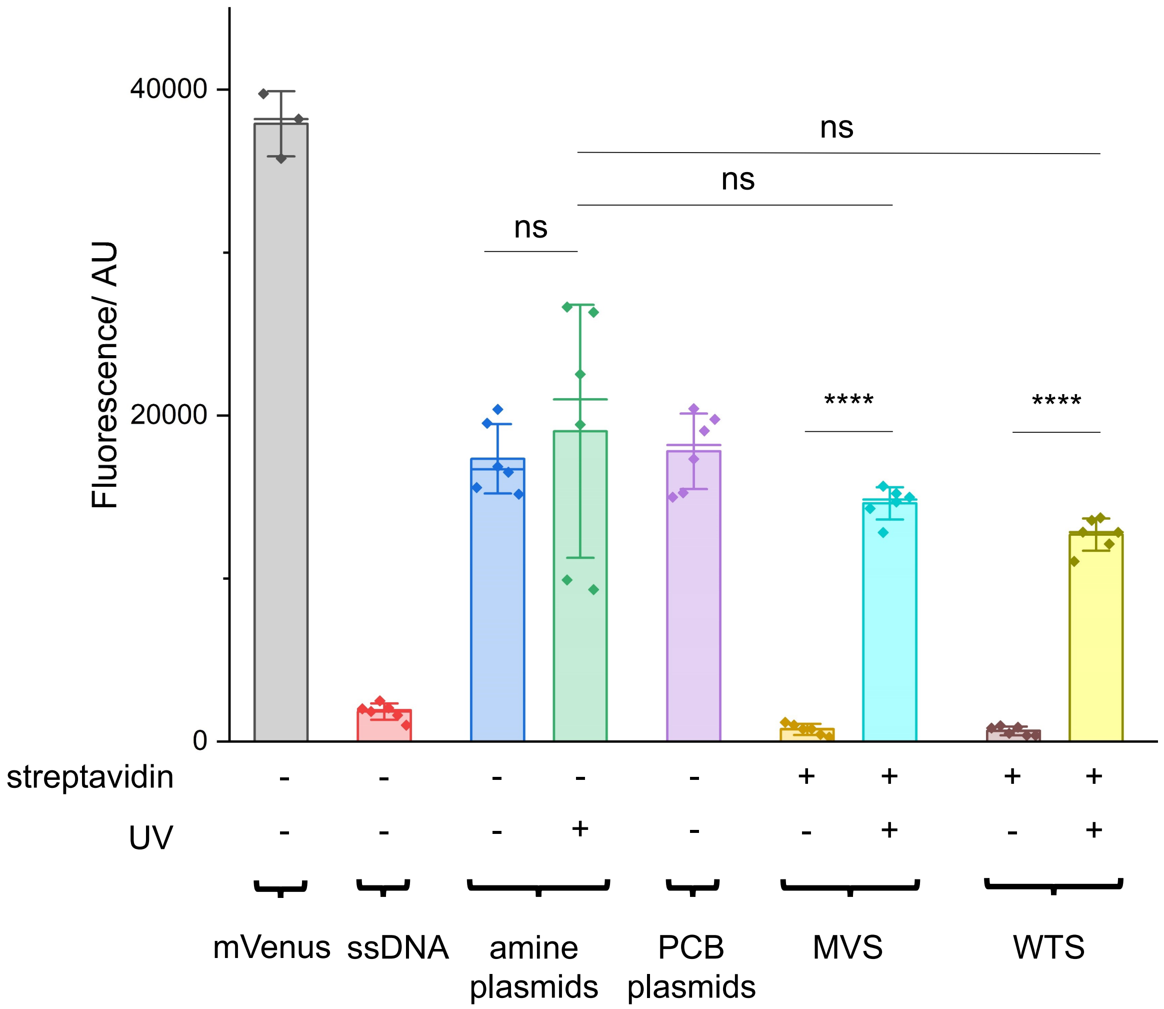

**SI Figure 8. CFPS of T7 NEEL plasmids including the native plasmid.** Plasmids containing the T7 promoter were prepared with NEEL and assessed using CFPS. Native plasmid and NTC were technical triplicates ($n$ = 3). Replicates, measurements, and statistical analysis were similar to those in **Figure 4B**.

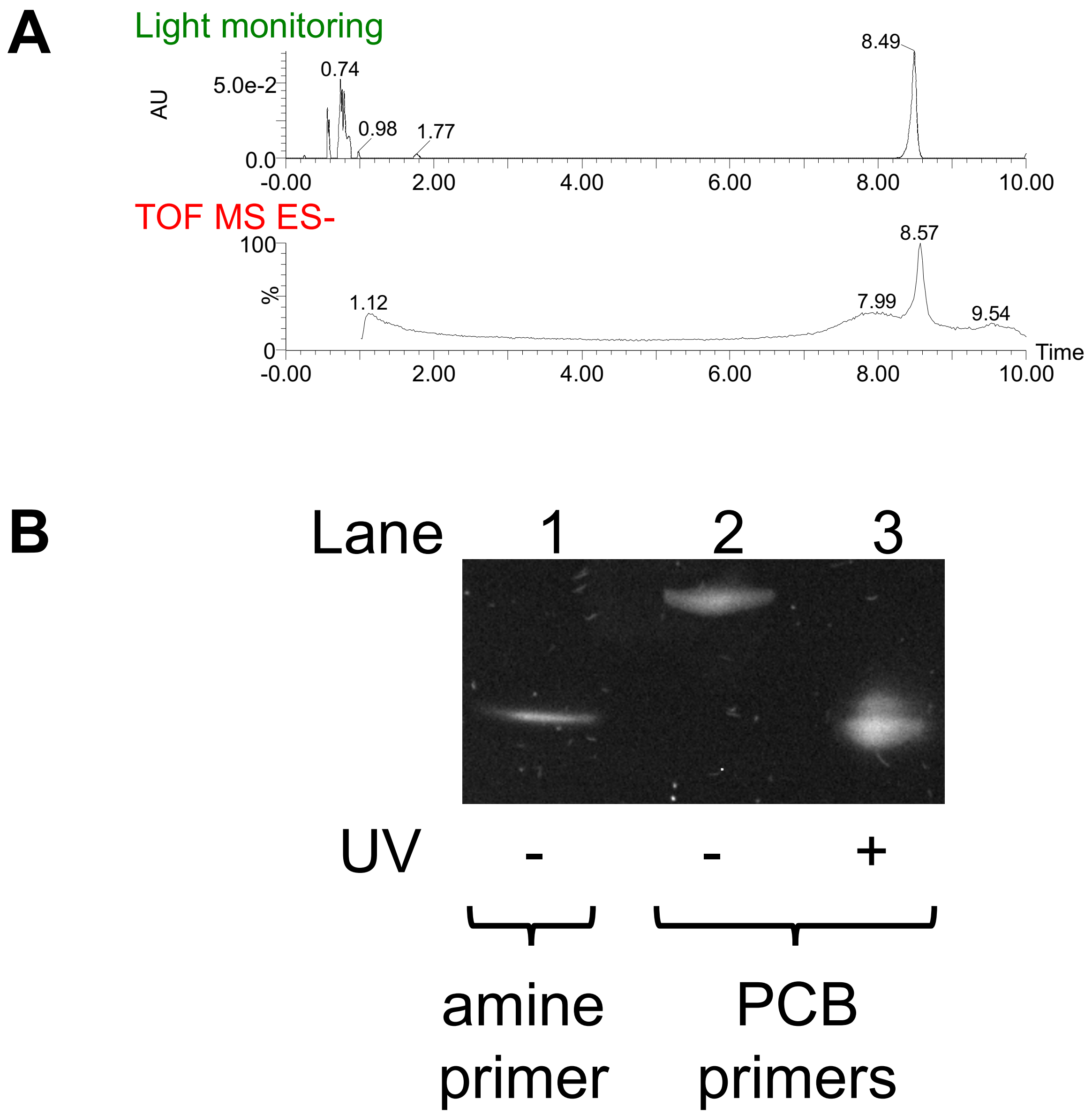

**SI Figure 9. Confirming synthesis of the CMV PCB oligo A.** LC-MS and **B.** Denaturing PAGE were used to confirm PCB attachment. MS gave a mass of 22076, which corresponded to the 9-PCBs oligo.

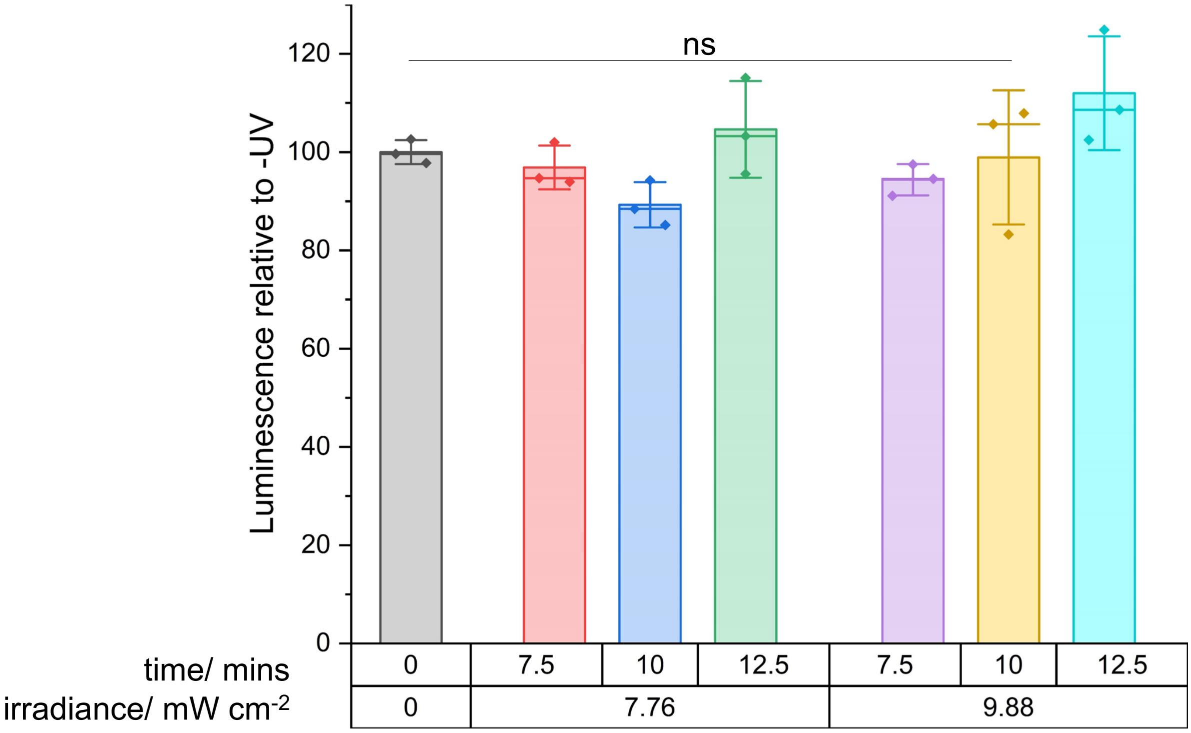

**SI Figure 10. Cell viability assay with optimised UV settings for transfection.** Cell viability assay with various UV settings. The conditions employed, 9.88 mW $cm^{-2}$ for 10 mins, had minimal damage to cells. Error bars showed standard deviations of technical triplicates ($n = 3$). Two-tailed, unpaired Student's t-test was performed. ns = non-significant.

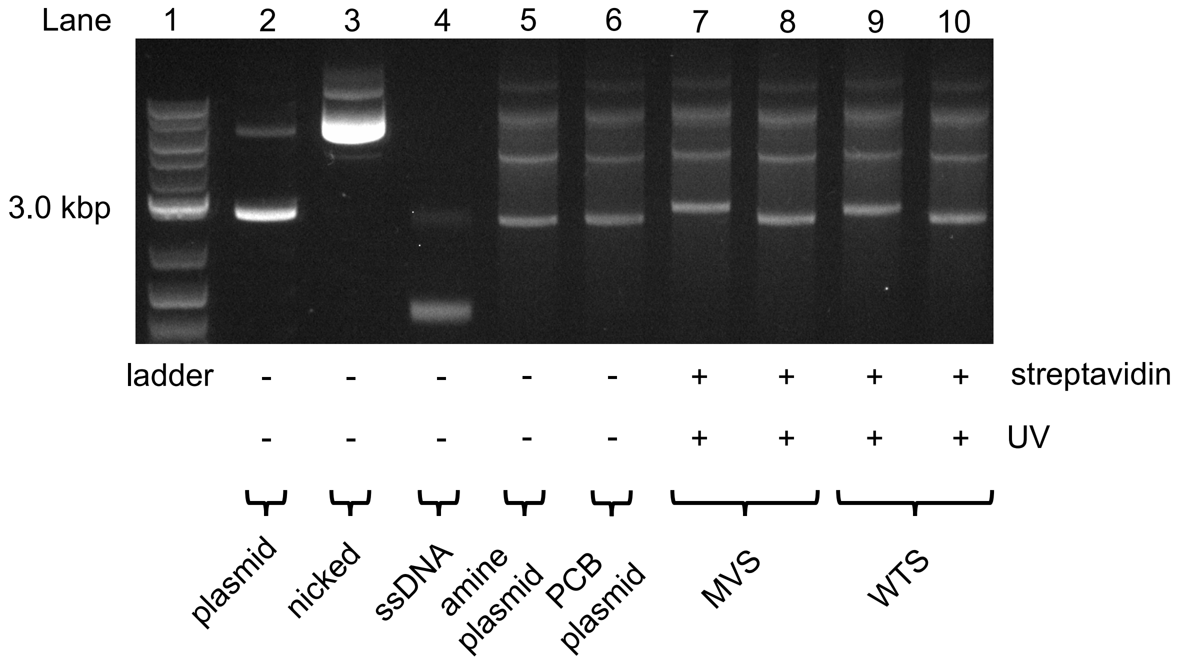

**SI Figure 11. CMV LA-plasmids and their uncaging.** The PCB plasmids were bound to streptavidins and uncaged using the optimised UV settings in **SI Figure 10**.

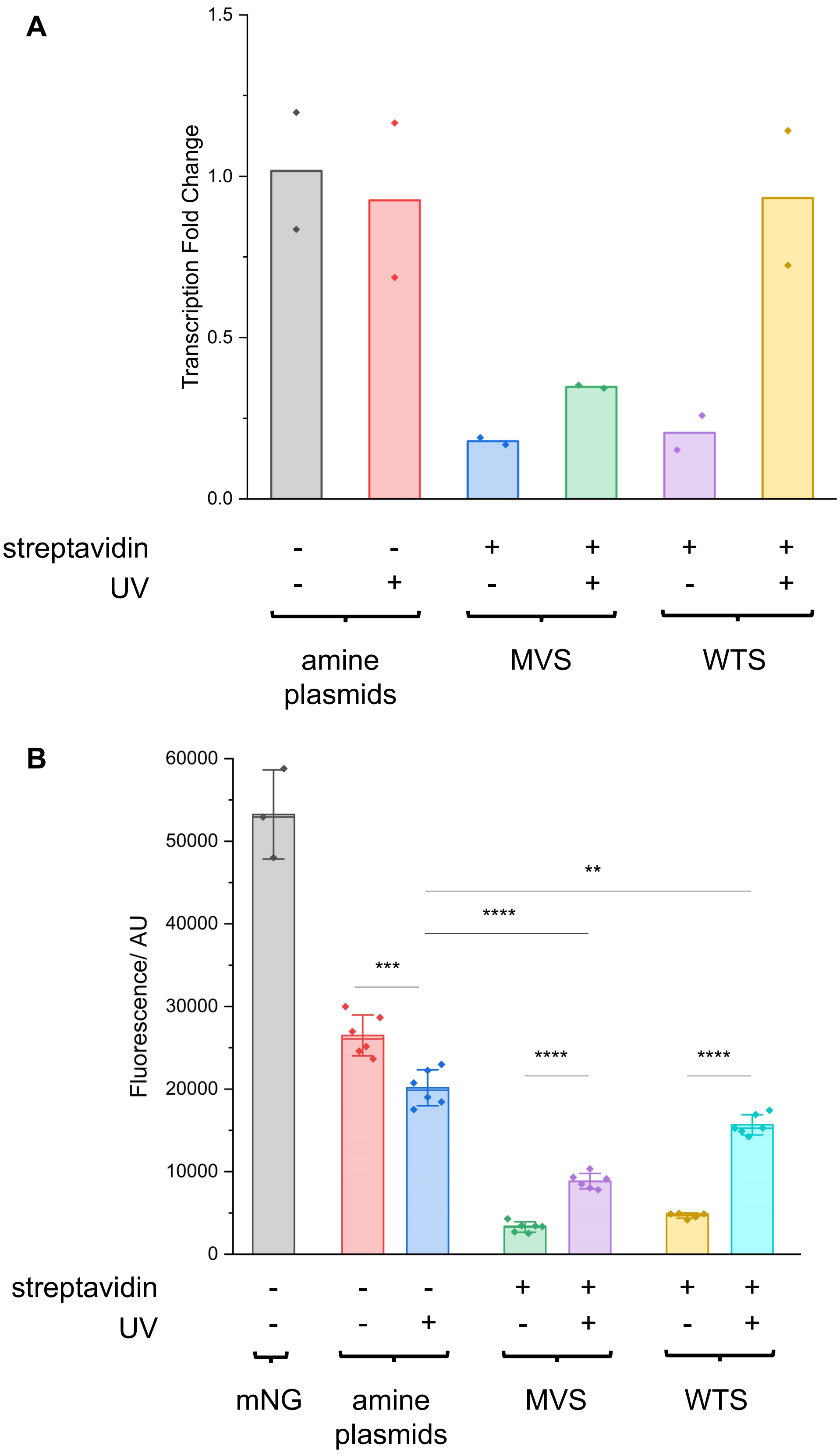

**SI Figure 12. RT-qPCR and fluorescence measurements of CMV NEEL plasmids, including the native plasmid.** Plasmids containing the CMV promoter were prepared with NEEL and assessed using **A.** RT-qPCR and **B.** Fluorescent plate reader. Replicates, measurements, and statistical analysis were similar to those in **Figure 5**.
